# *In situ* structure of bacterial 50S ribosomes at 3.0 Å resolution from vitreous sections

**DOI:** 10.1101/2025.06.03.657669

**Authors:** Ashraf Al Amoudi, Rozbeh Baradaran, Xukun Yuan, Fēi Wú, Andreas Naschberger

**Affiliations:** Electron Microscopy Lab, Imaging and Characterization Core Labs, King Abdullah University of Science and Technology (KAUST), 23955 Thuwal, Saudi Arabia; King Abdullah University of Science and Technology (KAUST), Biological and Environmental Science and Engineering Division, 23955 Thuwal, Saudi Arabia; Structural Biology STP, The Francis Crick Institute, 1 Midland Road, London NW1 1AT, United Kingdom

## Abstract

*In situ* high-resolution structure determination is limited to samples thin enough to be penetrated by the electron beam during imaging. Sample thinning involves focused ion or plasma beam milling of specimens to produce lamellae with thicknesses as low as 100-150 nm. However, surface damage caused by the milling process can extend 30-60 nm deep, restricting the usable lamella thickness. This imposes limitations on single-particle analysis of macromolecular complexes due to elevated structural noise, which cannot be avoided *in situ* because of the dense cellular environment. Alternative methods capable of producing thinner samples are needed to reduce background. Here, we demonstrate that high-resolution structures at side-chain level, free of orientation bias, can be obtained from vitreous sections prepared by cryo-ultramicrotomy, both *in vitro* and *in situ*. We optimized the method to produce sections as thin as ∼40 nm, free from significant surface damage. Using this approach, we determined the structure of the 50S ribosomal subunit *in vitro* at 2.8 Å and *in situ* at 3 Å from bacterial cells. These results lay the foundation for future *in situ* studies of smaller complexes using CEMOVIS, as well as for methodological advances aimed at achieving compression-free sectioning.

## Introduction

*In situ* cryo-electron microscopy (cryo-EM) enables the high-resolution structural determination of macromolecular targets in their near-native, hydrated state, embedded within the preserved cellular ultrastructure ^1^. Typically using cryo-Electron Tomography (cryo-ET), cryo-fixed cells or tissues are imaged by acquiring multiple tilt series, which are subsequently computationally aligned and processed to generate tomograms. These tomograms are representing reconstructed volumes corresponding to a portion of a cell ^2^. Using subtomogram averaging, the particles of interest within the volume are localized and extracted as subvolumes, which can then be aligned and averaged to obtain high-resolution reconstructions *in situ* ^3^. However, due to the crowded cellular environment and the poor contrast and signal-to-noise ratio (SNR) caused by relatively thick samples and structured background noise, high-resolution *in situ* structures are currently restricted mainly to large molecular assemblies, such as ribosomes ^4–7^.

More recently, it has been demonstrated that *in situ* cryo-EM is not limited to tomography alone but also *in situ* single-particle analysis (SPA) is applicable on cellular lamellas ^8–11^. This was first shown by resolving the photosystem I-II supercomplex of red algae *in situ* at an overall resolution of 3.2 Å ^8^. A similar approach was later applied to determine structure of the respiratory chain supercomplexes at 2.8 Å resolution (locally up to 1.8 Å) in unthinned mitochondria isolated from porcine tissue ^10^. To robustly identify particles in the crowded cellular environment using 2D projections of small cellular volumes for *in situ* SPA, several two-dimensional template matching (2DTM) algorithms have been developed ^12–15^. Those programs account for the background noise of overlapping densities from other molecules by applying either whitening filters ^12,15^ or a frequency-dependent signal-to-noise scoring function ^13^. 2DTM not only provides the x- and y-coordinates of the target molecule in the micrograph but also estimates the Euler angles and the z-position by searching across different contrast transfer function (CTF) values. Three software packages have been developed for this purpose and are available as free tools, including GiSPA ^16^, cisTEM ^15^, and HRTM ^12^.

The advantage of *in situ* single-particle EM over tomography lies in its significantly higher throughput, with acquisition times on the order of seconds rather than the several minutes required for tomographic tilt-series collection ^17^. However, the downside is that the entire content of the respective lamella is projected into 2D. Unlike tomography, there is no opportunity to computationally remove structural noise from regions above and below the target molecule to enhance the contrast of the specific tomographic z-slice. Hence, although *in situ* single-particle cryo-EM remains a viable alternative to tomography, the significantly higher background noise from cellular components in 2D imaging limits contrast and SNR, consequently, the ability to resolve molecules at high resolution, generally restricting analysis to large molecular assemblies^16^.

For *in situ* data collections, the sample must be thin enough to allow sufficient penetration of the electron beam through the tissue or cellular sample, while at the same time minimizing the damage to the biological material caused by inelastic scattering ^18^. In practice, the sample thickness for *in situ* studies is usually kept around the inelastic mean free path of the electrons, which depends on the accelerating voltage (e.g., 200 nm at 120 kV and ∼350 nm at 300 kV) ^19,20^. Typically, samples between 100 and 200 nm in thickness are prepared and should not exceed 500 nm. Several strategies were applied to achieve this: (i) Using cells or viruses that are thin enough to image directly (e.g., *Mycobacteria* with widths of 0.2–0.5 µm) ^21,22^, (ii) Imaging the cell cortex or the leading edge (lamellipodia of mouse embryonic fibroblasts) ^12^, which is typically thin enough to be penetrated by the beam (0.1–0.5 µm) , (iii) Focused ion beam (FIB) milling to thin the sample to a desired thickness (0.1–0.5 µm) to generate a thin lamella suitable for data collection ^23–25^. The FIB-milling approach is particularly powerful as it is not limited to specific cell or tissue types, enabling the study of a wide range of biological specimens. However, FIB milling also has several major drawbacks. First, the volume of specimens located above and below the chosen imaging section within the cells will be irreversibly removed and cannot be used for additional data collection of potentially interesting areas in those parts of the cell **(Fig. 1a)**. In addition, the radiation damage caused by the ion or plasma beam during removal of the biological material is substantial and can penetrate as deep as 30–60 nm from the surface of the lamella ^18,26^. This means that, for instance, in a 100 nm thick lamella, optimally only about 40 nm of the biological material at the center remains undamaged, whereas in the worst-case scenario, the entire volume of the lamella exhibits significant radiation damage. Furthermore, the detection of fluorescence-labeled proteins using Cryogenic Correlative Light and Electron Microscopy (Cryo-CLEM) in the z-direction remains challenging for FIB-milled lamellae, as identification of the z-plane in which the target resides is not very accurate **(Fig. 1a)** ^27,28^. Additionally, although recent technical advancements using plasma-based FIB scanning electron microscopes (FIB-SEM) which improved milling speeds substantially ^29^, FIB milling remains a relatively low throughput method ^1^. Automated FIB lamella generation typically yields several tens of lamellae per day ^30^, which is also limited by the buildup of ice contamination from residual water molecules in the vacuum column on the freshly prepared lamellae over time ^31^. Finally, the thickness of FIB-generated lamellae has a lower limit of about 100 nm ^18,26^. Going thinner than this leads to ion beam-induced damage within the lamella, leaving basically no undamaged region that could be imaged. However, to determine high resolution structures of smaller proteins, thinner lamellae are required to achieve sufficiently good contrast and SNR ^7,16^.

**Figure 1:**
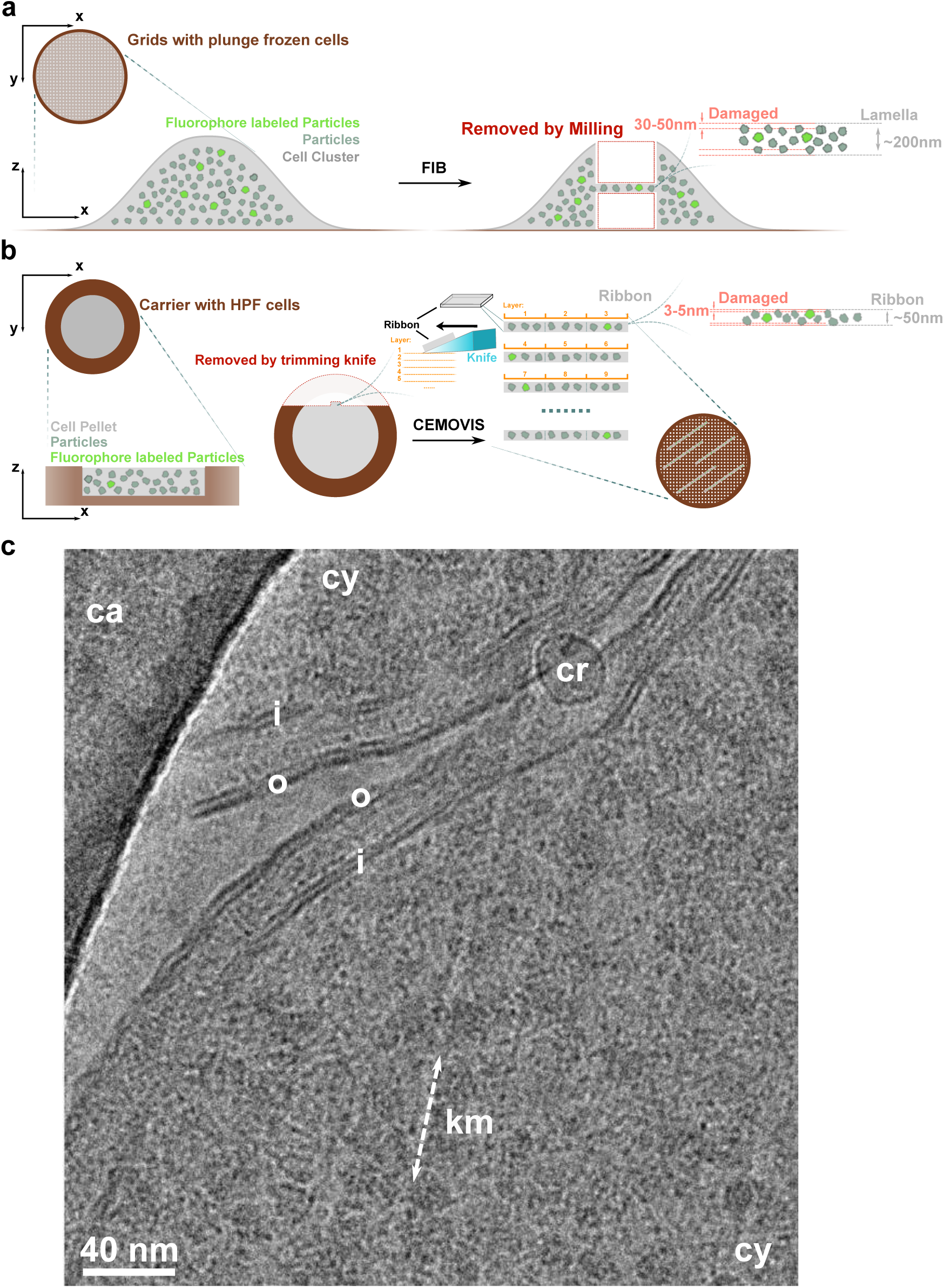
Using CEMOVIS to cut optimized ultrathin sections suitable for high-resolution data collection. **a**, FIB lamella production irreversibly destroys the material in the *z*-direction above and below the lamella with loss of fluorophore-labeled targets. The radiation damage was estimated to extend up to 60nm. **b,** Vitreous sections in the *z*-direction remain connected, forming a ribbon that can be attached to the grid, thus avoiding the loss of potentially useful fluorophore-labeled targets for imaging. The surface damage in CEMOVIS sections is typically restricted to a thinner surface layer. **c**, Representative crevasses-free vitreous section from *E. coli* cells imaged at high magnification shows high electron transparency suitable for high-resolution data collection (km: knife marks, cy: cytosol, o: outer membrane, i: inner membrane, cr: crystalline ice, and ca: carbon).

An alternative approach for producing thin sample of vitrified biological material is the use of cryo-electron microscopy on vitreous sections (CEMOVIS)^32–35^. The technique uses high-pressure freezing (HPF) to cryofix a sample ^36–38^, typically 100-200 µm thick, followed by slicing it into ultrathin sections using diamond knives ^39^. This overcomes most of the previously described limitations found for FIB-milled samples. CEMOVIS can cut sections as thin as 25-30 nm (nominal thickness also known as feed), which is within the same thickness range as the ice layers used in plunge-frozen samples for SPA ^40^. Surface damage caused by knife marks depends on the knife edge radius *r_k_* (3.2-4.4 nm +/- 0.5 nm) and is believed to be limited to 3-5 nm on each side of the section ^41^. Importantly, the method preserves the structural continuity in the z-direction, as all sections remain physically connected and are laid out sequentially like a film strip (ribbon) onto the EM grid **(Fig. 1b)**. This is especially useful for detection of fluorescence-labeled proteins in cryo-CLEM, as it preserves adjacent sections in the z-direction. Because each section remains attached to the next, the z-axis information is effectively projected into two dimensions **(Fig. 1b)** ^42^. The positional accuracy in the z-direction is determined by the section thickness. Furthermore, a skilled operator can achieve high throughput with CEMOVIS, generating several hundred low-artifact sections within a few hours of cryo-trimming and sectioning, with the closed-system design that minimizes ice contamination ^32^. However, even the best sections will inevitably suffer from knife marks on the surface and, more critically, from compression artifacts in the cutting direction (up to 60%) ^43^. This compression becomes increasingly severe as the section thickness decreases and may dampen all high-resolution signals of any macromolecules present in the cellular ultrastructure ^43,44^. While most studies using CEMOVIS have been carried out at low resolution ^42,45,46^ a recent study reported a 3.5 Å resolution structure of the 60S ribosome from *Saccharomyces* obtained from 100–150 nm thick sections ^47^. Here, we report a 3.0 Å *in situ* structure of the bacterial 50S ribosomal large subunit from ∼40nm thick sections, demonstrating that structural determination is possible using thinner high-quality vitreous sections from *Escherichia coli* (*E. coli)* cells. In addition, we show that this technique can also be applied to purified macromolecules, offering a potential solution to the preferred orientation problem frequently encountered in single-particle plunge-frozen samples. Finally, we discuss potential sources of compression artifacts and propose future directions aimed at eliminating or minimizing these, with the goal of establishing CEMOVIS as a viable alternative for *in situ* cryo-EM studies.

## Results

### Cutting ultrathin vitreous sections from *E. coli* cells optimized for SPA

The schematic in **Supplementary Figure 1** summarizes the various types of damage that can occur during the cutting of vitrified biological material using CEMOVIS ^43^. However, if performed under optimal conditions, all cutting artefacts can be eliminated or minimized, except for surface knife marks and compression caused by the cutting force. For example, crevasses can be avoided by cutting sections thinner than 50 nm ^43^. In HPF for CEMOVIS, dextran (20-40% w/v) is commonly added as a cryoprotectant to achieve sample vitrification ^34^. Dextran also facilitates smoother sectioning due to its plastic-like properties after vitrification. Since we aimed for SPA, we used a very dense suspension of *E. coli* cells to ensure their presence in as many acquisition areas as possible (**Supplementary Figure 2a)**. This allowed us to reduce the dextran concentration to 10%, as the high cell density displaced water between the cells, thereby reducing the need for cryoprotectant. Following vitrification of the *E. coli* suspension, the sample was sectioned using a cryo-ultramicrotome. Sections were cut as thin as possible, with nominal thicknesses (feed) ranging from 30 to 50 nm. They were attached to conventional copper EM grids (Quantifoil 1.2/1.3, Cu 300) by electrostatic charging **(Supplementary Figure 2b)** ^48^. We observed that the smaller 1.2 µm holes stabilized the sections more effectively than the larger 2 µm holes, likely due to their narrower geometrical constraints and hence better mechanical support. Additionally, we found it crucial to attach the sections to a grid surface free of contamination. Any residual ice present on the grid prior to section attachment led to poor adhesion and high drift during data collection. No additional modifications, such as carbon or other types of coatings, were required to stably attach the sections to the grids. The process yielded approximately 130 sections per 30 minutes, demonstrating the high-throughput capabilities of the method **(Supplementary Video 1)**. Inspection of the vitreous sections at high magnification (165,000×) showed high quality, electron transparent, crevasse-free, sections of *E. coli* cells, suitable for data collection using SPA **(Fig. 1c)**.

### SPA of purified 50S ribosomal subunit at 2.8 Å resolution from vitreous sections without orientation bias

The obtained high-quality sections from cells prompted us to investigate whether high-resolution structure determination could be achieved, first in a simplified *in vitro* system. Our reasoning was that if purified ribosomes were high-pressure frozen and sliced into ultrathin sections for imaging, their orientations in the microscope should be entirely random. This is in contrast to plunge freezing, where the air-water interface and thin ice layer often lead to preferred orientations ^49^ **(Supplementary Figure 3a,b)**.

We prepared highly concentrated bacterial ribosomes to aim HPF followed by sectioning (**Supplementary Figure 3c)**. Previous reports indicated that single-particle structures can, in principle, be determined in the presence of cryoprotectants ^50^. However, we removed dextran entirely to achieve the best possible contrast and SNR. We reasoned that the high concentration of isolated ribosomes would itself act as an effective cryoprotectant and enable full vitrification upon HPF. Ribosomes were cryofixed without the use of cryoprotectant using HPF, however, we employed a customized 25 µm carrier (Wohlwend GmbH, Sennwald, Switzerland) to minimize the sample volume and ensure full vitrification. There were no signs indicating significant crystalline ice formation, demonstrating that the sample was fully vitrified **(Fig. 2a)**. Motion correction and CTF estimation were performed using cryoSPARC ^51^ and revealed usable signal up to a resolution of 3.3 Å **(Supplementary Figure 3d,e)**. The data was processed in cryoSPARC **(Supplementary Figure 5, Table 1)**. We used template particle picking followed by particle extraction and 2D classification. However, the initially obtained 2D class averages were not successful **(Supplementary Figure 4)**, presumably due to the presence of numerous damaged particles due to the knife (see below in next section) and the expected random angular distribution with all viewing angles present, which likely resulted in too few particles per angle per class. To overcome this, a 50S ribosome map reconstructed from the same sample under conventional plunge-freezing conditions (in buffer) was used as a 3D reference in heterogeneous refinement, together with four junk classes **(Supplementary Figure 5, 6a, Table 1)**. Several iterative rounds of 3D classification and refinement using this reference allowed enrichment of ribosomal particles. Using those resulted in a final reconstruction of the 50S ribosomal subunit at 2.8 Å resolution **(Supplementary Figure 6b, Table 1)**.

**Figure 2:**
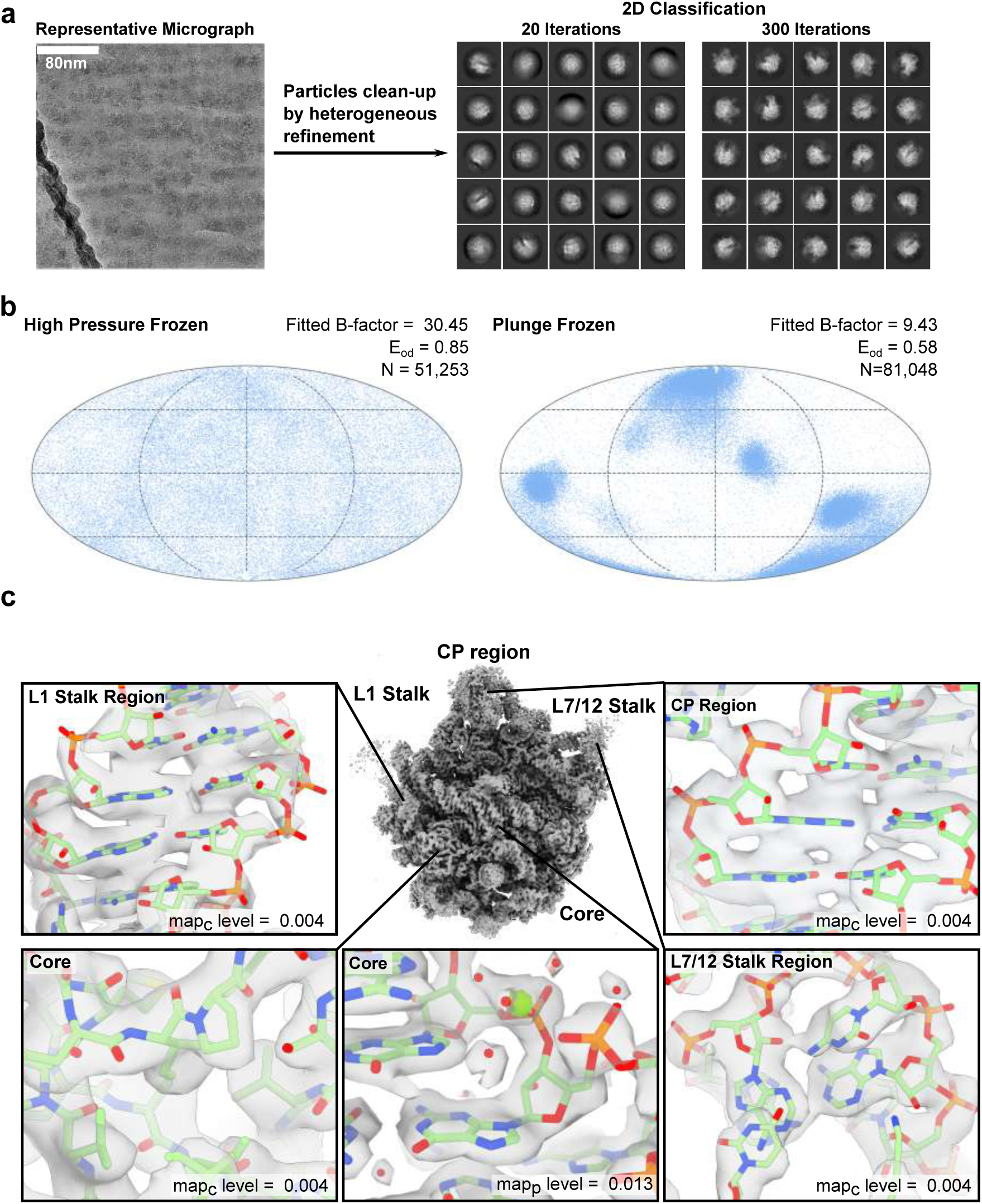
SPA of purified ribosomes at 2.8 Å from vitreous sections. The white scale bar corresponds to 80 nm. **a,** Representative micrograph of purified ribosomes from vitreous sections. After initial 3D cleanup by heterogeneous refinement, 2D classes can be obtained using 300 iterations, whereas in standard settings (20 iterations) the classes do not converge. **b,** Angular distribution plot comparison analyzed with cryoEF. The HPF sample shows no preferred orientation, whereas the plunge-frozen sample exhibits strong preferred angles. Several metrics are shown: estimated B-factor, orientation-distribution efficiency (Eod), and particle count. **c,** 2.8 Å resolution map of purified ribosomes from vitreous sections shown as an isosurface representation. Map quality is illustrated in five close-up views from the 50S ribosomal subunit core (rRNA and protein), the central protuberance (CP), the L7/L12 stalk region, and the L1 stalk region. The map levels of the consensus map (map_c_) and the sharpened post-processed map (map_p_) are indicated.

**Table 1.**
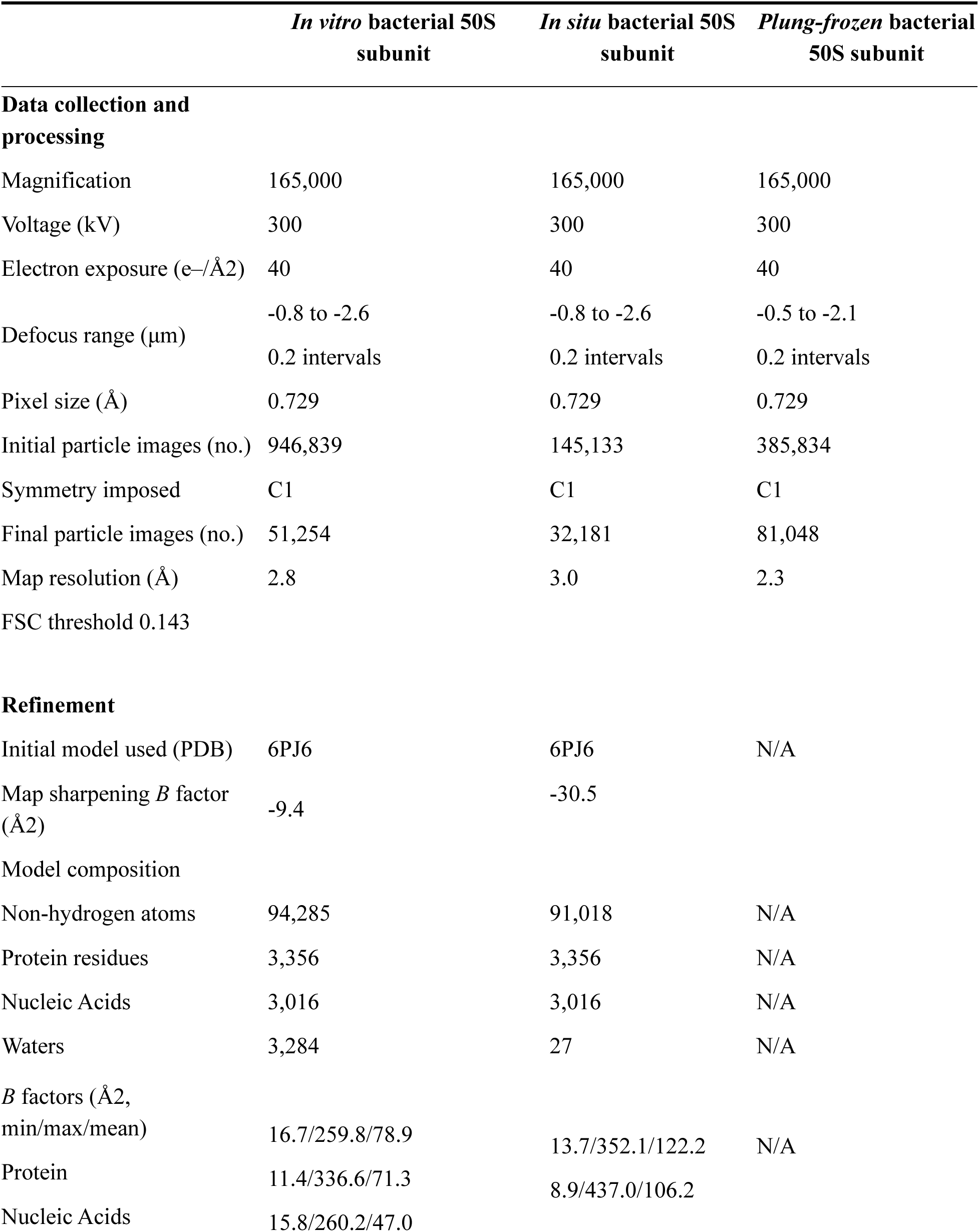
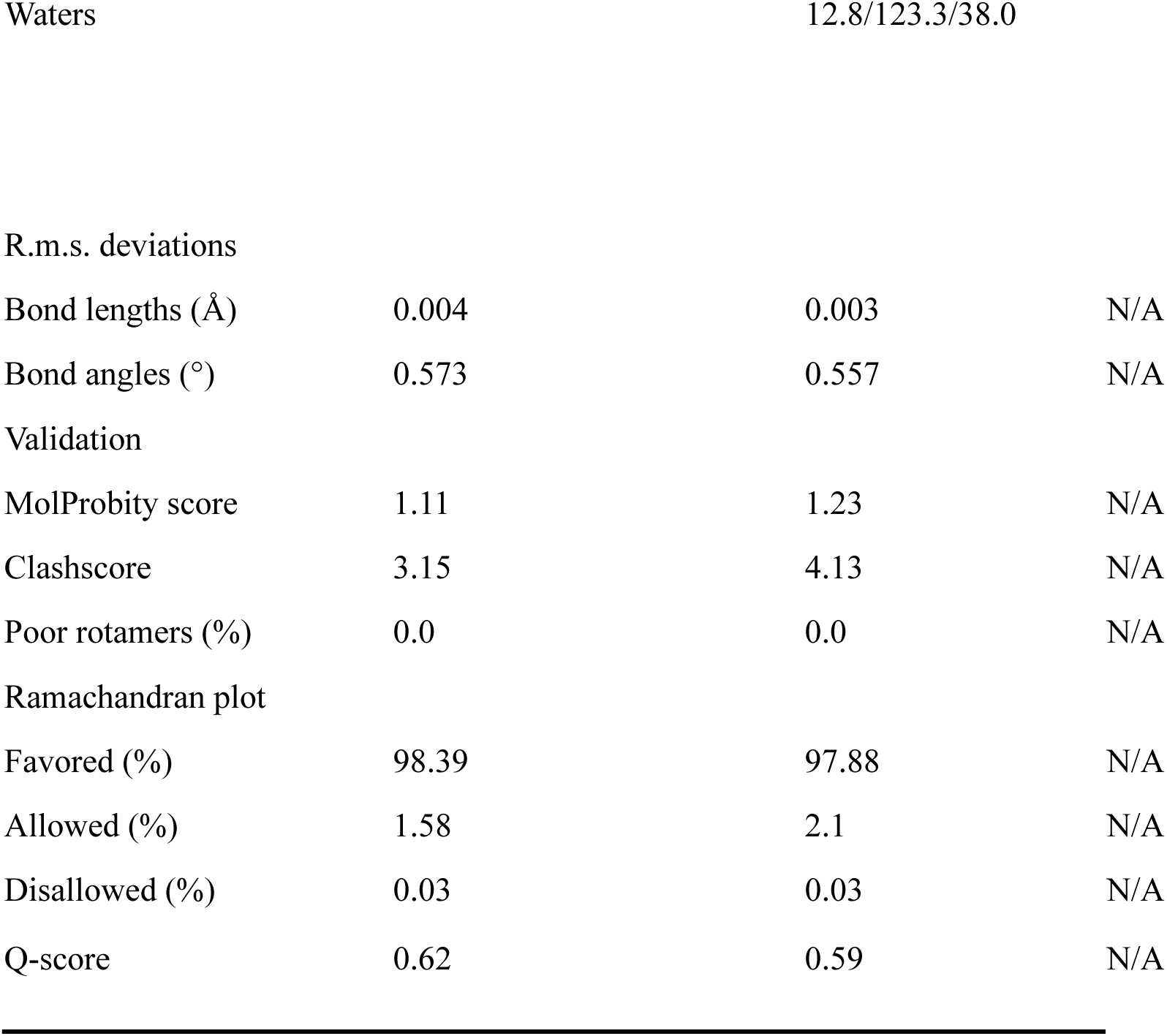
Summary of data collection, refinement, and validation statistics for the *in vitro* and *in situ* bacterial 50S ribosome structures obtained from vitreous sections.

It is worth noting that this cleaning step also works with maps downloaded from the PDB when used as references for heterogeneous refinement. We also noticed that 2D classification became effective when using a cleaner set of particles after the initial 3D heterogeneous refinement **(Fig. 2a)**. Although, the classes remained blurry with the default settings, increasing the number of iterations to 300 yielded clear 2D class averages across a wide range of views **(Fig. 2a)**. Comparison of the angular distribution with a plunge-frozen reference dataset showed that orientation bias was absent **(Fig. 2b)**. The obtained map showed side-chain level resolution across different regions of the ribosome, and clear density for water molecules in the 50S ribosomal core **(Fig. 2c)**. Further, the *in vitro* reconstruction from vitreous sections shows no significant damage or differences compared to the plunge-frozen data, suggesting that compression does not significantly affect the macromolecular structure **(Supplementary Figure 7a-c)**.

In conclusion, we demonstrate that structure determination at 2.8 Å resolution is possible using purified proteins from vitreous sections, and that this approach can overcome the orientation bias problem. Early 3D-based processing using an initial reference is essential, as 3D classification efficiently enriches intact particles when many are broken, whereas 2D classification fails to converge.

### Knife-induced damage in ultrathin sections limits the size of molecules that can be structurally preserved

In the dataset, we were also able to detect 70S monosomes. However, we observed that the 30S subunit and the central protuberance (CP) region of the monosome appeared less well resolved than expected, suggesting that some form of damage may have occurred. Size measurements of the low-pass filtered map from the plunge-frozen monosome indicated that the intact part of a vitreous section must be at least 28 nm thick to accommodate the full ribosome, including more flexible regions such as the L7/L12 stalk **(Fig. 3a)**. To estimate the thickness of our sections, we collected 18 tilt series from *in situ*-generated vitreous sections, reconstructed tomograms for each of them, and measured the section thickness at multiple sampling points **(Fig. 3b)**. A mean section thickness *t* of 48.36 ± 3.74nm was estimated **(Fig. 3c)**. This value reflects the actual section thickness after compression has occurred. However, the critical parameter that determines whether an object was damaged during the cutting process due to the object size is not this compressed thickness *t*, but the nominal feed value, denoted as *f*, set at the microtome i.e. before compression has occured. The relationship between the feed thickness *f* and the actual section thickness *t* is given by the Equation (2) **(Supplementary Figure 1)**. The cutting angle in our study was set to 41° (35°+6° clearance) and can therefore be considered as constant and the feed *f* was chosen between 30-50 nm. Additionally, knife marks typically reduce the usable thickness by about 5 nm from each side ^41^, leaving only a ∼30 nm undamaged central zone in the section **(Fig. 3d)**. This model allows us to calculate the probability that a spherical object will end up within this undamaged zone. The expected probability can be estimated using Equation (3) **(Fig. 3d)** and is only 15% for the 50S subunit and 5% for 70S monosomes **(Fig. 3e-g)**. Further theoretical considerations show that even if the feed is increased to 50 nm, the point at which crevasses begin to form, the probability of 70S ribosomes ending up undamaged in the central region still remains low (25%) **(Fig. 3g)**. Thus, cutting thicker than 50 nm would introduce surface damage in the form of crevasses ^43^, which may in turn damage particles located beneath them. Therefore, ultrathin sections (∼40nm) are necessary to avoid crevasses and are best suited for smaller molecular targets (e.g., <50S subunit).

**Figure 3:**
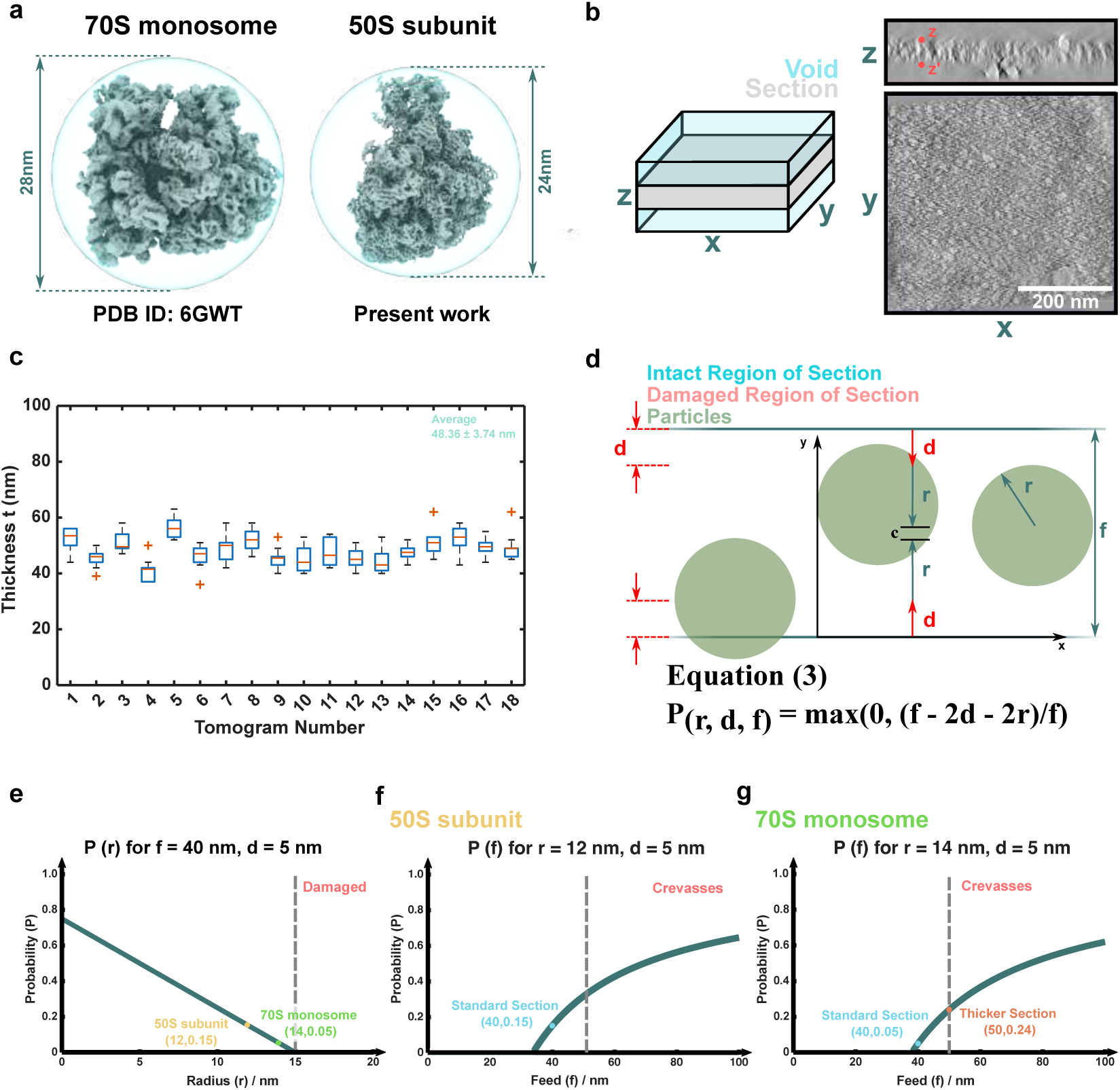
Size limitations in cryo-EM of vitreous sections. **a,** Diameter of bacterial 70S and 50S ribosomal subunits. **b,** Representative tomogram of vitreous sections with indicated measurement points used for section thickness estimation. The white scale bar corresponds to 50 nm. **c,** Measured thicknesses *t* after compression from 18 tomograms is shown in box plot representation. For each tomogram, 10 locations were sampled (n = 180). The box plots show the 25th–75th percentiles (blue box), median (red line), interquartile range (black whiskers), and outliers (red crosses). The mean section thickness was 48.36 ± 3.74 nm (cyan line) (see also Supplementary Data 1). **d,** Simplified statistical model for particle damage along the z-direction. Sections are randomly sliced with feed *f*, and particles are assumed to be spheres of radius *r*. Knife marks of depth *d* create damaged regions at the section surfaces, leaving an intact central region. The probability of a particle remaining undamaged is given by Equation (3) as the fraction of allowed center positions *c* within *f*. Overlaps in x-y are neglected. **e,** Probability from Equation (3) plotted versus particle radius (*f* = 40 nm, *d* = 5 nm). Undamaged particles occur with ∼5% probability for monosomes (*r* = 14 nm) and ∼15% for 50S subunits (*r* = 12 nm). **f-g,** Equation 3 plotted for *r* = 12 and 14 nm and *d* = 5 nm as a function of *f*. Intact particle probability increases with *f*, but crevasses appear above 50 nm.

### *In situ* 3.0 Å reconstruction of the 50S ribosomal subunit using SPA and vitreous sections from bacterial cells

We next used the vitreous sections prepared in **Fig. 1** and collected data for SPA. The motion correction and CTF estimation for the *in situ* data **(Supplementary Figure 3a,b)** showed fitting comparable to the *in vitro* dataset **(Supplementary Figure 8a,b)**. Again, we started with 3D heterogeneous refinement using the map obtained from the plunge-frozen dataset as a reference. Of note, similar to the *in vitro* case, only after an initial round of 3D classification by heterogenous refinement, stable 2D class averages were obtained by increasing the number of iterations to 300 **(Fig. 4a)**. Several rounds of 3D classification were required to separate the heterogeneous particles into a homogeneous subset. This heterogeneity likely reflects a combination of the higher structural variability within the cell and physical damage from the sectioning process. The number of particles restricted the number of computational parameters to achieve stable classifications, and only a small subset of particles remained after several rounds.

**Figure 4:**
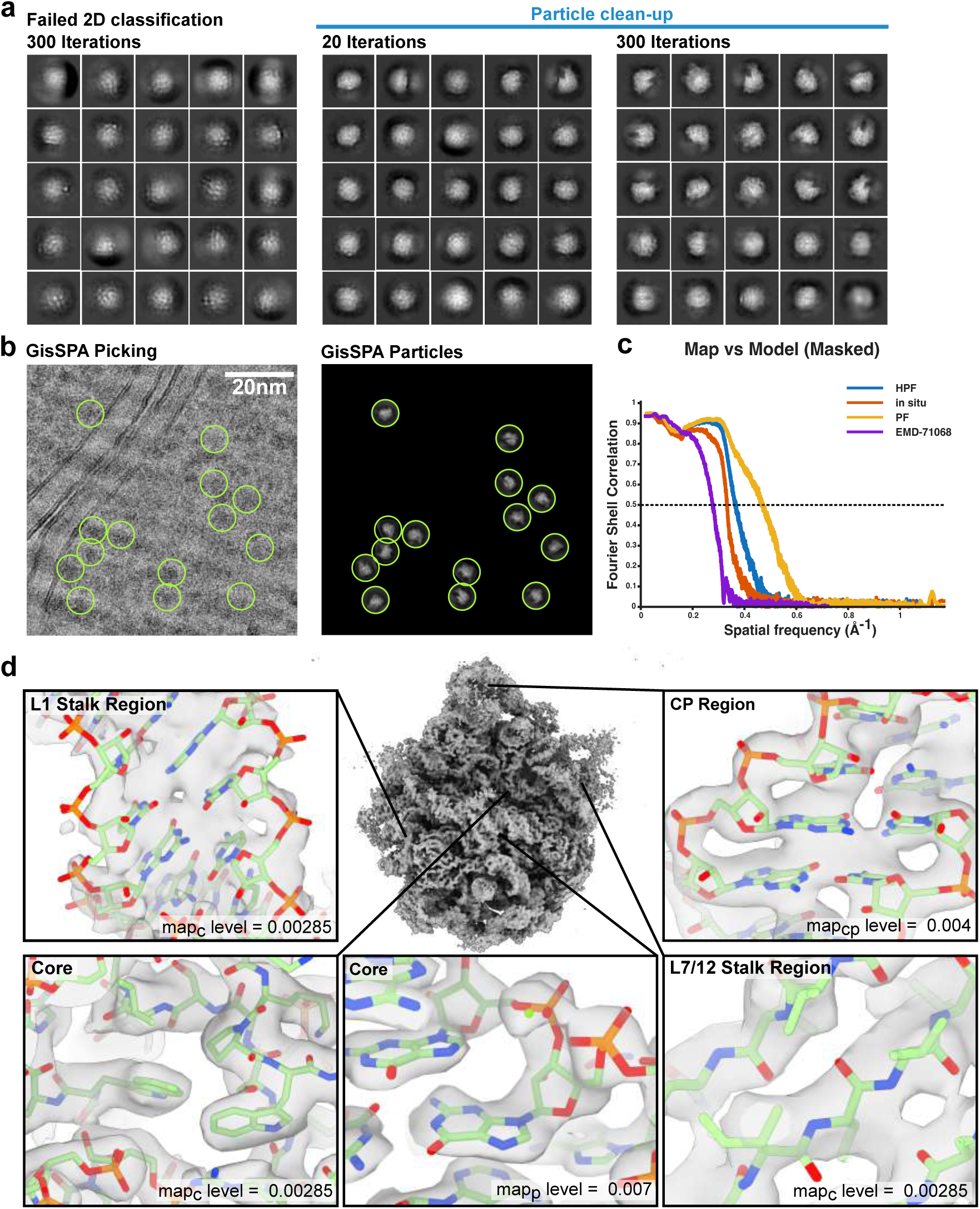
*In situ* cryo-EM structure of the bacterial 50S ribosomal subunit at 3.0 Å resolution from vitreous sections. **a,** Initial 2D class averages (left), and after one round of heterogeneous refinement using 20 and 300 iterations (right). **b,** 2DTM using GisSPA software with the plunge-frozen 3D reference volume of the 50S subunit as the search model. The panel on the right shows the matched templates projected with the correct orientation with the background entirely removed. The white scale bar corresponds to 20 nm. **c,** Map-to-model FSC comparing the *in situ* CEMOVIS and *in vitro* CEMOVIS datasets, the plunge-frozen dataset, and previous CEMOVIS dataset (EMD-71068 with PDB: 6PJ6 ^68^). **d,** High-resolution map showing side-chain details of the 50S ribosomal subunit with positions and map levels indicated (map_p_: postprocessed/sharpened map, map_c_: consensus/unsharpened, map_cp_: masked refined map on CP).

To increase the number of particles identified from the micrographs, we performed GisSPA 2DTM ^16^ on the motion-corrected micrographs using the 50S ribosomal subunit from the plunge-frozen reference as the search model **(Supplementary Figure 5)**. We were able to match 122k ribosomes **(Fig. 4b, Supplementary Figure 9a)**, from which, after several rounds of classification, a reconstruction at 3.0 Å resolution was obtained using 32,181 particles in RELION ^52^ **(Fig. 4c, Supplementary Figure 9, Table 1)**. The map shows high-resolution features at the side-chain level **(Fig. 4d)**. The CP region and some rRNA segments on the ribosomal surface were flexible, as indicated by the decreased resolution in these regions **(Fig. 4d, Supplementary Figure 9b)**. To show that the CP region, although more flexible, is still intact, we performed focused refinement using a smaller mask centered on the CP region **(Fig. 4d, Supplementary Figure 9)**. The analysis shows that the density exhibits the typical expected improvement, indicating that the internal integrity of the CP is preserved. We further validated our approach by repeating the 2DTM analysis using a search model derived from the Protein Data Bank (PDB ID: 6PJ6), in which density for two ribosomal subunits (L27 and L22) were omitted prior to 2DTM **(Supplementary Figure 10a)**. The resulting 3D reconstruction displayed well-resolved density for the two omitted subunits, confirming that template bias was limited **(Supplementary Figure 10a,b)**. Finally, to analyze the SNR of each obtained dataset, we calculated the mean SNR per particle as a function of resolution **(Supplementary Figure 10c, Supplementary Data 2)**. The analysis shows that while the SNR of the plunge-frozen dataset is comparable to that of the *in vitro* data, the SNR in the *in situ* dataset is decreased, most likely due to the higher structural noise.

In summary, we obtained a high-resolution reconstruction of the *E. coli* 50S ribosome *in situ* using vitreous sections and SPA, showing that CEMOVIS is a valid alternative method to study macromolecules *in situ* at 3 Å resolution.

## Discussion

In this work, we demonstrate using single-particle cryo-EM that high-resolution structure determination from vitreous sections at 3 Å resolution is possible for both isolated and *in situ* 50S ribosomes. Two recent studies similarly demonstrate the potential of CEMOVIS for high-resolution structural studies ^47,53^. The first study used CEMOVIS to produce 150–200 nm thick sections of high-pressure frozen lysozyme nano crystals as a novel approach in electron diffraction experiments ^53^. The authors concluded that the main advantage of the method is its high throughput of protein structural determination by electron diffraction, enabled by the large number of crystals that can be produced using CEMOVIS and the reduced background noise resulting from the removal of surrounding solvent through the sectioning process. The more recent study showed that *in situ* determination of the 60S ribosomal subunit in yeast cells is feasible using SPA applied to 100 nm-thick sections prepared by CEMOVIS ^47^. The report demonstrates an overall resolution of 3.5 Å, showing that even with greater section thickness, high resolution is achievable. Thicker sections also experience less compression and increase the likelihood of obtaining undamaged particles not affected by the knife **(Fig. 3)**. However, in sections thicker than 70 nm (which corresponds to a feed of 50 nm), crevasses are unavoidable. Crevasses are alternating patterns of fissures and compressed material on the section surface, creating dark areas unsuitable for data collection. The fissures can also damage the biological material in their vicinity.

Unlike our present work, the authors reported problems with the attachment of their sections to the grids. We have summarized our success rates in **Supplementary Table 1**. For example, in the *in situ* case, we are able to attach the section in about 80% of the attempts, and 50% of micrographs with the best quality were used. Their poor attachment can be explained by the greater thickness, which leads to crevasses and consequently, poor section attachment. Moreover, thicker sections possess greater mass, which in our experience significantly compromises their stable attachment. This is likely due to the insufficient effect of electrostatic charging to adhere the vitreous sections to the support film of the EM grid. Hence, while additional methodological effort is required for thicker sections, coating or other grid modifications are not needed for stable attachment of thinner sections.

We optimized CEMOVIS for sections as thin as ∼40 nm (thickness *t*), which improves the SNR and enables high-resolution structure determination. When sections are cut under optimal conditions, most artifacts can be minimized or completely avoided, except for compression and knife marks in the cutting direction. The knife marks in CEMOVIS are believed to be only 3-5 nm in extent ^41^, which is significantly less than the surface damage typically seen in FIB-milled lamellae (30-60 nm) ^18,26^. While compression remains a concern for large-scale deformations of the cellular environment, the overall structural integrity at the macromolecular level appears to be preserved. This is supported by our successful reconstruction of intact 50S ribosomal particles in sections of isolated ribosomes and *in situ*. Nevertheless, further efforts are needed to enable a compression-free sectioning process that fully preserves the cellular environment, to establish CEMOVIS as a robust accepted method not only for high-resolution structure determination but also for analysis of the overall cellular architecture. We propose two promising directions to address the compression problem. First, as described by Equation (1) in **Supplementary Figure 1**, compression decreases with the cutting angle α. A decrease in compression with decreasing knife angle was also shown experimentally ^54,55^. This implies that at a knife angle of 0°, no compression would occur. Currently, however, knives are too fragile to use at angles lower than 25°. Future development to improve knives may help overcome this limitation. Second, we have previously shown that compression-free vitreous sections can be produced using an oscillating knife which virtually reduces the cutting angle down to 0° ^44,56^. Although the reproducibility was insufficient, the method was not extensively optimized, and technical advancements in oscillating knife technology over the last two decades may have improved the precision of the process. These technical developments will not only enable the production of compression-free vitreous sections but also eliminate crevasses in thicker sections, thereby extending the applicability of CEMOVIS to large macromolecular complexes.

The major advantage of CEMOVIS lies in its ability to produce much thinner sections with relative small surface deformation and significantly higher throughput compared to FIB-milled lamellae. Such thin sections are not achievable with current FIB-milling techniques, as FIB introduces damage up to 60 nm from each side, limiting lamella thickness to 100–150 nm. Nevertheless, we observed in our study that 70S monosomes suffer from degradation, as indicated by the low resolution densities of more flexible elements such as the CP and the 30S small subunit. Several scenarios could explain the source of these lower resolution densities. One possibility is that HPF might compromise the molecular integrity. However, a recent study demonstrated that intact 70S monosomes can be obtained from high-pressure frozen samples, ruling out this explanation ^30^. That study achieved a ribosome reconstruction at ∼4 Å resolution, confirming structural preservation. Another possibility is that compression during sectioning contributes to these lower resolution densities at the macromolecular level. In our study, the 50S large subunit remained intact, making it unlikely that compression alone accounts for the observed phenotype *in vitro*. Importantly, monosomes show lower resolution densities specifically in fragile regions, such as the CP region of the 50S subunit, which are generally flexible and likely the first regions to be affected. Yet, the same region appears preserved in the 50S ribosomal subunit *in vitro*, further indicating that compression is not the primary cause of monosome degradation. We therefore conclude that the monosome is simply too large to fit within the undamaged zone of 30 nm-thick sections. Our analysis, shown in **Fig. 3**, supports this idea. Nevertheless, since the *in situ* environment differs from that of isolated samples, the findings from *in vitro* data cannot be directly extrapolated to the *in situ* context. Our *in situ* reconstruction shows increased flexibility at the surface compared to the *in vitro* map. This could reflect the larger conformational space expected *in situ* relative to *in vitro*. Alternatively, compression *in situ* may have led to greater apparent structural flexibility, for example in the CP region. Another possible explanation is that, due to the low number of particles per micrograph, the fraction of undamaged particles was insufficient to allow clear separation of knife-damaged particles during classification. Collecting larger datasets might increase the number of intact particles, thereby allowing more reliable separation from the knife-damaged ones. To obtain high-resolution 70S monosome reconstructions, thicker sections would be required to accommodate entire monosomes within the undamaged interior of the sections. However, there is a size limit, as sections thicker than 50 nm typically develop crevasses, leading to additional sources of damage. Finally, future work should clarify whether high-resolution reconstructions can also be obtained for molecules with a molecular weight below 0.5 MDa.

In conclusion, we demonstrate that CEMOVIS is a valid technique for high-resolution structure determination at the macromolecular scale, both *in situ* and *in vitro*. Future efforts will focus on overcoming compression artifacts to establish CEMOVIS as a widely adopted method for *in situ* structural biology both at the ultrastructural and macromolecular levels.

## Methods

### *E. coli* culture and preparation of a highly concentrated cell suspension suitable for HPF

*E. coli* cells (BL21) were plated onto conventional LB agar plates and grown overnight at 37 °C. The next day, a single colony was picked and inoculated into a 50 ml Falcon tube containing 20 ml of LB medium. This starter culture was grown overnight at 37 °C and 220 rpm in a shaking incubator. On the following day, three 1 l flasks, each containing 200 ml of LB medium, were inoculated with 1 ml, 5 ml, and 10 ml of the overnight culture, respectively, and incubated at 37 °C and 220 rpm. OD₆₀₀ was monitored continuously at 30-minute intervals. Once one culture reached an OD₆₀₀ of 0.4, it was harvested by centrifugation for 10 minutes at 4,000 × g (Thermo Scientific Sorvall LYNX 6000 Superspeed Centrifuge, Fiberlite F14-6 x 250y Fixed Angle Rotor) at 4 °C. The remaining two cultures were discarded and had only been grown as backups in case the culture initiated with a 10 mL inoculum exceeded the intended harvesting point at OD₆₀₀ = 0.4. The resulting pellet was resuspended in 10 ml of 1× PBS and centrifuged again at 4,000 × g for 10 minutes at 4 °C (Eppendorf Centrifuge 5910 Ri , Eppendorf Rotor S-4xuniversal). The final pellet (5 ml) was carefully resuspended in 5 ml of 20% dextran (prepared in 1× PBS). From this suspension, 1 ml was transferred into a 1.5 ml Eppendorf tube and centrifuged for 10 minutes at 4 °C at 4,000 × g (Eppendorf Centrifuge 5425 R, Eppendorf FA-24x2 Fixed Angle Rotor). The supernatant was removed, and the resulting paste-like pellet was directly used for high-pressure freezing.

### Purification of ribosomes from *E. coli* cells

To purify ribosomes from *E. coli* cells, we followed the protocol described by Yi Cui *et al.,* 2022 ^57^. In brief, the LB culture described in the previous section was grown to an OD₆₀₀ of ∼0.9. The culture was harvested by centrifugation at 4,000 × g (Thermo Scientific Sorvall LYNX 6000 Superspeed Centrifuge, Fiberlite F14-6 x 250y Fixed Angle Rotor) for 10 minutes at 4 °C. The resulting 2.7 g pellet was carefully resuspended by pipetting up and down in a 50 ml Falcon tube using 12 ml of Buffer A (50 mM Tris-acetate, pH 7.7; 60 mM potassium glutamate; 14 mM magnesium acetate). Then, 700 µl of 5 mg/ml lysozyme (prepared in Buffer A) were added, and the suspension was incubated for 30 minutes at 37 °C with shaking at 700 rpm (Eppendorf ThermoMixer C). The cell suspension was then subjected to freeze-thaw cycles in liquid nitrogen to lyse the cells and release the lysate. Subsequently, 20 µl of DNase I (Invitrogen, 18047-019) were added to degrade genomic DNA, and the mixture was incubated for 1 hour at 4 °C. The lysate was cleared by centrifugation at 10,000 rpm for 1 hour at 4 °C (Thermo Scientific Sorvall LYNX 6000 Superspeed Centrifuge, Fiberlite F14-6 x 250y Fixed Angle Rotor). The resulting supernatant (∼20 ml) was filtered through a 0.22 µm filter to remove residual debris. The filtrate was then diluted with 20 ml of Buffer B (20 mM Tris-HCl, pH 7.7; 500 mM ammonium chloride; 10 mM magnesium acetate; 0.5 mM EDTA; 30% sucrose; 7 mM β-mercaptoethanol). This mixture was centrifuged at 170,000 × g (average) for 1.5 hours at 4 °C (Thermo Scientific Sorval WX+ Ultracentrifuge SeriesSureSpin™, 632 Swing-out Ultracentrifuge Rotor). The supernatant was discarded, and the pelleted ribosomes were washed with 500 µl of Buffer C (20 mM Tris-HCl, pH 7.7; 6 mM magnesium acetate; 30 mM potassium chloride; 7 mM β-mercaptoethanol). Finally, the pellet was resuspended in 50 µl of Buffer C. The RNA concentration of the purified ribosomes was estimated to be 34 µg/µl using a NanoDrop spectrophotometer.

### Grid preparation and data acquisition of plunge-frozen *E. coli* ribosomes

For the reference dataset, we used the same purified sample as for the isolated ribosomes in vitreous sections. Quantifoil R2/2 copper grids with a 2 nm carbon support layer and 300 mesh size (100 pieces, Jena Bioscience) were glow-discharged using a GloQube Plus instrument (Quorum Technologies, UK) for 30 seconds at 35 mA. The purified bacterial ribosomes were centrifuged at maximum speed (benchtop centrifuge) for 10 minutes at 4 °C to pellet aggregated proteins. From the resulting supernatant, 3 µl was applied onto the Quantifoil grids and incubated for 30 sec in the 100% humidity chamber of a Vitrobot Mark IV (FEI Company, USA). The grid was blotted (blot force = 0) for 3 sec and subsequently plunge-frozen in liquid ethane. Grids were clipped and loaded into a 300-kV Titan Krios G4 microscope (Thermo Fisher Scientific, EPU v3.5.1.6034 software) equipped with a Selectris Energy filter (Slit width 10 eV) and a Falcon 4i direct electron detector (Thermo Fisher Scientific). Grids were screened for quality control based on particle distribution and density, and images from the best grids were recorded. Micrographs were recorded at a nominal magnification of 165,000x, corresponding to a calibrated pixel size of 0.729 Å. The dose rate was 8 electron per physical pixels per second, and images were recorded for 3 s in the EER format, corresponding to a total dose of 40 e/Å^2^. Defocus range was set between −0.5 μm and −2.1 μm (0.2 μm intervals). Gain-corrected image data were acquired.

### Cryo-EM data processing of plunge-frozen *E. coli* ribosomes

Processing was started in cryoSPARC Live v4.6.0 ^51^ with motion correction, CTF estimation, and blob picking. A total of 385,834 particles were picked and extracted using a 192-pixel box size (Fourier-cropped from 500 pixels) and used for 2D classification. The high quality 2D class averages were subjected to *ab initio* job with 4 classes as input. The class with the highest-resolution features (the best class) was refined to 3.9 Å, and this map was then used as input for heterogeneous refinement with two good classes and three junk classes, using all 385,834 picked particles as input. The best class contained 124,933 particles and was refined to a resolution of 3.01 Å. This particle set was also exported to RELION v5.0 ^52,58^ and re-extracted without binning using a 500-pixel box. After refinement followed by classification into 4 classes (T = 25, E = 6) , the class corresponding to the 50S ribosomal subunit was used for a final refinement, which resulted in a map at 2.3 Å resolution (81,048 particles). Of note, the choice of T values is empirical and must be tested by trial and error. The standard T = 4 is not always optimal.

### High-Pressure Freezing (HPF)

Two types of samples were prepared for high-pressure freezing: purified *E. coli* ribosomes and dense *E. coli* cell pellets. Purified ribosome suspensions were loaded into custom-fabricated 25 µm-deep membrane carriers (Wohlwend GmbH, Sennwald, Switzerland) without any cryoprotectant. The ribosome was concentrated to 34 ug/ul which was sufficient to ensure efficient heat transfer and achieve vitrification upon rapid freezing. Dense *E. coli* cell pellets were prepared as described above. These cell pellets were then loaded into standard 100 µm-deep membrane carriers (Leica Microsystems). The concentration is an empirical parameter and must be determined by trial and error, increasing it until vitrification without crystal formation is possible. The absence of Bragg diffraction rings in diffraction mode confirmed that the sample was vitrified.

All samples were vitrified using an EMPACT II high-pressure freezer (Leica Microsystems) at ∼2100 bar and cooling rates exceeding 20,000 °C/s. Immediately after freezing, carriers were transferred into pre-cooled metal storage boxes under liquid nitrogen to maintain vitrified conditions. Sample carriers were stored in liquid nitrogen until further processing. HPF conditions, including pressurization rates and cooling curves, were monitored using the EMPACT II’s onboard diagnostic tools to ensure reproducibility across preparations.

### Cryo-Ultramicrotomy and CEMOVIS

Frozen carriers were mounted under liquid nitrogen into a Leica EM UC7/FC7 cryo-ultramicrotome (Leica Microsystems). Prior to sectioning, sample faces were trimmed using a cryo-trimming knife to expose the vitrified material and to produce a sample block with a side ∼ 100 µm for cryosectioning. Ultrathin vitreous sections with a feed ranging from 30–50 nm (in 80% of all sections, a feed of 40 nm was used during cutting) were cut at –140 °C using a non-oscillating 35° CEMOVIS or cryo-immuno diamond knife (Diatome Ltd., Nidau, Switzerland), with a cutting speed of 0.5–1.0 mm/s. An additional 6° angle is introduced by the clearance angle of the knife, which is required to leave space between the knife and the upper surface of the next section. This results in a total cutting angle of *α* = 41° (35° + 6°).

The compression is given by (see also Supplementary Figure 1):

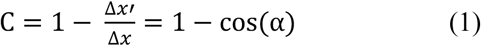

Where α is the cutting angle and Δx represents a volume element of the sample in the feed direction that is deformed into a compressed volume element Δx’ after sectioning. The relationship between the nominal section thickness (also known as the feed, *f*) and the compressed section thickness *t* is given by:

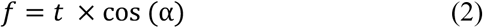

Sections were collected onto 300 mesh copper Quantifoil R1.2/1.3 grids (Quantifoil Micro Tools GmbH) using a double micromanipulator system, ensuring minimal contact and mechanical stress. Section gliding on the knife surface was made smooth by using the electrostatic discharging with the ionizer. Vitreous sections were attached to EM grids by brief electrostatic charging, with the grids charged immediately before section retrieval using the built-in ionizer (Charging Device, Leica Microsystems).The ionizer in the cryo-microtome was used as follows: (i) A ribbon of vitreous cryo-sections, which acts as an electrical insulator, is guided over a grounded, conductive EM grid. (ii) The ionizer’s electrode, positioned approximately 10-20 mm from the grid, is switched to a “charge mode.” This mode applies a negative DC high voltage to the electrode, creating a strong negative field. (iii) This strong negative field causes negatively charged ions from the ionized atmosphere to accumulate on the surface of the non-conductive vitreous ribbon. (iv) An electrostatic force of attraction is thereby generated between the negatively charged ribbon and the grounded, conductive EM grid, pulling the sections down securely onto the gird.

Each grid was visually inspected under the microtome stereomicroscope to assess section quality, integrity and good attachment. Grids were stored in a firmly closed cryo-grid boxes inside the cryo-ultramicrotome and maintained at cryogenic temperatures throughout transport to the cryo-electron microscope for subsequent cryo-electron tomography or single-particle cryo-electron microscopy.

### Data collection and SPA processing of purified ribosomes from vitreous sections

The state of vitreous water was verified using the electron diffraction.

Data were collected on a Titan Krios G4 microscope (Thermo Fisher Scientific) equipped with a Falcon 4i camera and a Selectris energy filter (slit width: 10 eV) from vitreous sections of purified ribosomes using EPU software (v3.5.1.6034) at a magnification of 165,000×, corresponding to a pixel size of 0.73 Å/pixel. A total of 8,435 movies were acquired with a total dose of 40 e⁻/Å² and a defocus range of –0.7 μm to – 2.6 μm (0.2 μm intervals).

**Supplementary Figure 3** illustrates the data processing workflow for isolated *E. coli* ribosomes prepared by HPF and CEMOVIS. The following pre-processing steps were performed with cryoSPARC Live v4.6.0 ^51^. Movie stacks were motion-corrected and dose-weighted using MotionCor2 v2.1.1 ^59^. Contrast transfer function (CTF) estimates for the motion-corrected micrographs were calculated with CTFFIND4 v4.1.13 ^60^. Particles were autopicked using 44 of the representative 2D classes averages containing different orientations as templates prepared from the plunge-freezing data. This resulted in 946,839 picked particles from 8,435 micrographs. Each particle was extracted at a box size of 96 pixels (5.2x binned). Subsequent image processing was carried out with cryoSPARC v4.6.0. An initial model generated from the plunge-freezing data was used as a reference volume to clean up the particles. Particles were classified by three rounds of three-dimensional (3D) heterogeneous refinement using 5 classes. Particles were then re-extracted at 500 pixels (unbinned) and an initial model was generated. Analysis of the initial model identified only the 50S ribosomal subunit in the reconstruction. The initial model was subsequently used for homogeneous refinement followed by non-uniform (NU) refinement to get a reconstruction at 3.7 Å overall resolution with C1 symmetry. CTF refinement (without 4th order aberrations) followed by another round of NU refinement improved the quality of the reconstruction. Particles were converted into a STAR file using Pyem and imported into RELION v5.0 for further processing ^52,58^.

Particles were re-extracted at 500 pixels (unbinned) followed by 3D refinement using the cryoSPARC map as the starting model. CTF refinement (without 4th order aberrations) followed by another round of 3D refinement improved the quality of the reconstruction. Bayesian polishing was performed, and subsequent 3D refinement on the polished particles resulted in reconstruction at 3 Å overall resolution with C1 symmetry. Further CTF refinement (without 4th order aberrations), Bayesian Polishing and 3D refinement improved the quality of the reconstruction to 2.74 Å overall resolution with C1 symmetry. To remove the particles that have been damaged by the CEMOVIS preparation, 3D classification was performed (without alignment) on the particles using five classes and a T value of 4. A class with 45% of particles (51,328 particles) was identified. Subsequently, the particles from this class were subjected to 3D refinement, CTF refinement, 3D refinement, Bayesian polishing and 3D refinement to yield a reconstruction to 2.74 Å overall resolution with C1 symmetry. Analysis of the angular distribution of the refined particles showed no orientation bias in comparison to the particles from the plunge-freezing data, which suffer major orientation bias **(Fig. 2c)**.

In order to improve the density for the CP region of the 50S ribosomes, focused 3D classification (without alignment) was performed on the consensus map using three classes and a T value of 4 with a mask covering the CP region of the 50S ribosomal subunit. Local 3D refinement of a class containing 34,616 particles using the same mask covering the CP region improved the density in this region.

All final reconstructions were sharpened using RELION post-processing. The local resolution estimations in **Supplementary Figure 5** were performed using RELION.

### Tilt-series acquisition and cryo-electron tomography of bacterial vitreous sections

Tilt-series of bacterial vitreous sections were acquired on a Titan Krios G4 transmission electron microscope (Thermo Fisher Scientific) equipped with a Falcon 4i direct electron detector and a Selectris energy filter with a slit width of 10 eV. The tilt series were collected using Tomography 5 software (Thermo Fisher Scientific) at a nominal magnification of 105,000×, corresponding to a pixel size of 1.19 Å at the specimen level. Images were recorded with dose-symmetric mode over a tilt range of –60° to +60° with an angular increment of 2° or 3°, using an electron dose of approximately 2-3 e⁻/Å² per tilt image. Tilt-series alignment was performed in IMOD ^61^ using the Etomo interface with fiducial-less patch tracking alignment. Three-dimensional reconstructions were generated by weighted back-projection to obtain tomograms for subsequent analysis. The section thickness was quantified using the XYZ-plane view of the tomograms in IMOD. For each tomogram, twelve pairs of z-coordinates corresponding to the upper and lower surfaces of the section at identical x-positions were manually measured. Thickness statistics and visualization were performed in MATLAB R2025a.

### *In situ* data acquisition and SPA processing of 50S ribosomes from vitreous sections

Grids containing vitreous sections from bacterial cells were imaged using a total electron dose of 40 e⁻/Å². A magnification of 165,000× was used on a Titan Krios G4 microscope (Thermo Fisher Scientific) equipped with a Falcon 4i camera and a Selectris energy filter (slit width: 10 eV), corresponding to a calibrated pixel size of 0.729 Å. In EPU software (v3.5.1.6034), holes were detected automatically and curated manually. The defocus range was set between –0.7 μm and –2.6 μm (0.2 μm intervals). A total of 28,790 movies were recorded and subsequently gain-corrected. Movies were initially imported into cryoSPARC Live v4.6.0 for motion correction ^51^. Blob picking was used to identify particles across the micrographs, and 3D heterogeneous refinement was performed using the plunge-frozen 50S ribosome reconstruction as the reference. However, after several rounds of classification and refinement, it became evident that too few particles remained to obtain an intact 50S ribosomal subunit at side-chain-level resolution. Therefore, we adopted a different processing strategy, as outlined in **Supplementary Figure 9.** The movies were imported into RELION 5.0 ^52^, and motion correction was repeated using MotionCor2 ^59^. Particle identification was carried out using GisSPA 2DTM software ^16^ with the plunge-frozen 50S map as the search model and a score threshold of 6.8 (the reference was band-pass filtered 400 to 8 Å). This resulted in the detection of 145,133 particles, which were extracted in a 500-pixel box without binning. A 3D classification without alignment or mask was performed using 3 classes, a T-value of 4, and an E-value of –1. The best class (86%) contained approximately 120,000 particles, which were then subjected to several alternating rounds of CTF refinement and Bayesian polishing, resulting in a consensus reconstruction at 2.96 Å resolution. In a second processing branch, several classification steps were performed to remove damaged particles. The first classification focused on the CP region using a mask covering the entire CP (T = 10, E = 4). The best class (60.5%) was selected for further processing. A subsequent classification used a peripheral mask (2 classes, T = 10, E = 4), and the best class (81.5%) was used in a final classification round covering the entire 50S subunit (2 classes, T = 10, E = 4). The best class (80.6%) was selected from the polished dataset and refined, resulting in a 2.98 Å consensus map. Finally, a focused refinement was carried out using a smaller mask on the CP region to improve local resolution.

### Model building, refinement, and figures

The 50S ribosome structure published in the PDB under the accession number 6PJ6 was rigid-body fitted using Chimera ^62^ into the *in vitro* and *in situ* maps of the structures determined from vitreous sections. This was followed by flexible fitting using Coot v0.9.8.95 ^63^ with activated self-restraints. The model was then manually refined in real space using Coot to correct side-chain torsions and other local deviations from the original model. The models were subsequently imported into ISOLDE ^64^ to correct all rotamer outliers, Ramachandran outliers, and to improve clash scores. Final refinement was performed using real-space refinement in PHENIX ^65^, and validation was carried out using the PDB validation server. Model-to-map FSCs were calculated using PHENIX. Iterative refinement with manual model building in Coot was continued until the model quality and model-to-map fit were sufficiently high. All figures were prepared using Chimera X ^66^. The angular distribution and metrics shown in Fig. 2 was calculated with cryoEF ^67^. The probability of a particle ending up in the undamaged zone and not being cut into parts by the knife as calculated in Fig. 3 is given by:

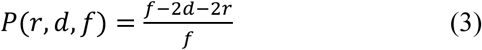

where *f* is the feed, *d* are the depth of the knife marks, and *r* is the radius of the particle. The mean SNR per particle in Supplementary Figure 10 was calculated using the following formula:

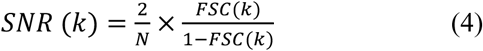

where FSC is the spatial frequency (*k*)-dependent cross-correlation between the half-maps, and *N* is the number of particles.

### Statistics and Reproducibility

In this manuscript, we determined the structure of the bacterial 50S ribosomal subunit using single-particle cryo-EM. All standard statistics are provided in Table 1, including the number of movies and particles, the gold-standard resolution estimate, and model validation and refinement metrics. Descriptive statistics for section thickness measurements are shown in Figure 3, where we report an *n* of 180 measured positions and present the data as a box plot including mean, standard error, and standard deviation. No statistical hypothesis testing was performed on these measurements.

## Data availability

The atomic coordinates of the bacterial ribosome structures obtained from vitreous sections were deposited in the Protein Data Bank under accession codes 9XFK (*in situ*) and 9XFL (*in vitro*). The corresponding density maps were deposited in the EMDB under accession codes EMD-66639 and EMD-66736, respectively. The maps of the CP region were deposited in the EMDB under accession codes EMD-66640 (*in situ*) and EMD-66749 (*in vitro*). The map from the plunge-frozen reference dataset was deposited in the EMDB under the accession code EMD-66841.

## Acknowledgments

This work was financed via the KAUST baseline fund. We thank the members of the IAC-EM Core Lab at KAUST, Lingyun Zhao, Alessandro Genovese and Rachid Sougrat for their continuous support with cryo-EM data collection and for maintaining the microscopes and associated equipment.

## Author Contributions Statement

AN and AA designed this study. AA prepared all vitreous sections used in this study. RB purified the ribosome sample and collected the *in vitro* data from vitreous sections. XY and RB prepared the cell suspensions for the *in situ* studies. AN, AA and RB collected and processed the *in situ* data. FW built and refined the atomic models. All authors jointly analyzed the data, prepared the figures, and wrote the manuscript.

## References

1 Noble, A. J. & de Marco, A. Cryo-focused ion beam for in situ structural biology: State of the art, challenges, and perspectives. Current Opinion in Structural Biology 87, 102864 (2024). 10.1016/j.sbi.2024.102864

2 Young, L. N. & Villa, E. Bringing Structure to Cell Biology with Cryo-Electron Tomography. Annu Rev Biophys 52, 573–595 (2023). 10.1146/annurev-biophys-111622-091327

3 Pyle, E. & Zanetti, G. Current data processing strategies for cryo-electron tomography and subtomogram averaging. Biochem J 478, 1827–1845 (2021). 10.1042/BCJ20200715

4 Tegunov, D., Xue, L., Dienemann, C., Cramer, P. & Mahamid, J. Multi-particle cryo-EM refinement with M visualizes ribosome-antibiotic complex at 3.5 A in cells. Nat Methods 18, 186–193 (2021). 10.1038/s41592-020-01054-7

5 Xing, H. et al. Translation dynamics in human cells visualized at high resolution reveal cancer drug action. Science 381, 70–75 (2023). 10.1126/science.adh1411

6 Zheng, W. et al. Visualizing the translation landscape in human cells at high resolution. bioRxiv (2024). 10.1101/2024.07.02.601723

7 Rickgauer, J., Choi, H., Lippincott-Schwartz, J. & Denk, W. Label-free single-instance protein detection in vitrified cells. (2020).

8 You, X. et al. In situ structure of the red algal phycobilisome-PSII-PSI-LHC megacomplex. Nature 616, 199–206 (2023). 10.1038/s41586-023-05831-0

9 Zhang, X., Xiao, Y., You, X., Sun, S. & Sui, S. F. In situ structural determination of cyanobacterial phycobilisome-PSII supercomplex by STAgSPA strategy. Nat Commun 15, 7201 (2024). 10.1038/s41467-024-51460-0

10 Zheng, W., Chai, P., Zhu, J. & Zhang, K. High-resolution in situ structures of mammalian respiratory supercomplexes. Nature 631, 232–239 (2024). 10.1038/s41586-024-07488-9

11 Lucas, B. A., Himes, B. A. & Grigorieff, N. (eLife Sciences Publications, Ltd, 2023).

12 Rickgauer, J. P., Choi, H., Moore, A. S., Denk, W. & Lippincott-Schwartz, J. Structural dynamics of human ribosomes in situ reconstructed by exhaustive high-resolution template matching. Mol Cell 84, 4912–4928 e4917 (2024). 10.1016/j.molcel.2024.11.003

13 Cheng, J., Li, B., Si, L. & Zhang, X. Determining structures in a native environment using single-particle cryoelectron microscopy images. Innovation (Camb) 2, 100166 (2021). 10.1016/j.xinn.2021.100166

14 Rickgauer, J. P., Grigorieff, N. & Denk, W. Single-protein detection in crowded molecular environments in cryo-EM images. Elife 6 (2017). 10.7554/eLife.25648

15 Lucas, B. A. et al. Locating macromolecular assemblies in cells by 2D template matching with cisTEM. Elife 10 (2021). 10.7554/eLife.68946

16 Cheng, J. et al. Determining protein structures in cellular lamella at pseudo-atomic resolution by GisSPA. Nat Commun 14, 1282 (2023). 10.1038/s41467-023-36175-y

17 Eisenstein, F. et al. Parallel cryo electron tomography on in situ lamellae. Nat Methods 20, 131–138 (2023). 10.1038/s41592-022-01690-1

18 Tuijtel, M. W. et al. Thinner is not always better: Optimizing cryo-lamellae for subtomogram averaging. Sci Adv 10, eadk6285 (2024). 10.1126/sciadv.adk6285

19 Grimm, R., Typke, D., Barmann, M. & Baumeister, W. Determination of the inelastic mean free path in ice by examination of tilted vesicles and automated most probable loss imaging. Ultramicroscopy 63, 169–179 (1996). 10.1016/0304-3991(96)00035-6

20 Diebolder, C. A., Koster, A. J. & Koning, R. I. Pushing the resolution limits in cryo electron tomography of biological structures. J Microsc 248, 1–5 (2012). 10.1111/j.1365-2818.2012.03627.x

21 Xue, L. et al. Visualizing translation dynamics at atomic detail inside a bacterial cell. Nature 610, 205–211 (2022). 10.1038/s41586-022-05255-2

22 Turonova, B. et al. In situ structural analysis of SARS-CoV-2 spike reveals flexibility mediated by three hinges. Science 370, 203–208 (2020). 10.1126/science.abd5223

23 Young, R. J., Dingle, T., Robinson, K. & Pugh, P. J. A. An application of scanned focused ion beam milling to studies on the internal morphology of small arthropods. Journal of Microscopy 172, 81–88 (1993). 10.1111/j.1365-2818.1993.tb03396.x

24 Marko, M., Hsieh, C., Schalek, R., Frank, J. & Mannella, C. Focused-ion-beam thinning of frozen-hydrated biological specimens for cryo-electron microscopy. Nat Methods 4, 215–217 (2007). 10.1038/nmeth1014

25 Rigort, A. et al. Focused ion beam micromachining of eukaryotic cells for cryoelectron tomography. Proc Natl Acad Sci U S A 109, 4449–4454 (2012). 10.1073/pnas.1201333109

26 Lucas, B. A. & Grigorieff, N. Quantification of gallium cryo-FIB milling damage in biological lamellae. Proc Natl Acad Sci U S A 120, e2301852120 (2023). 10.1073/pnas.2301852120

27 Li, S. et al. HOPE-SIM, a cryo-structured illumination fluorescence microscopy system for accurately targeted cryo-electron tomography. Commun Biol 6, 474 (2023). 10.1038/s42003-023-04850-x

28 Pierson, J. A., Yang, J. E. & Wright, E. R. Recent advances in correlative cryo-light and electron microscopy. Curr Opin Struct Biol 89, 102934 (2024). 10.1016/j.sbi.2024.102934

29 Berger, C. et al. Plasma FIB milling for the determination of structures in situ. Nat Commun 14, 629 (2023). 10.1038/s41467-023-36372-9

30 Berger, C., Watson, H., Naismith, J. H., Dumoux, M. & Grange, M. Xenon plasma focused ion beam lamella fabrication on high-pressure frozen specimens for structural cell biology. Nature Communications 16, 2286 (2025). 10.1038/s41467-025-57493-3

31 Tacke, S. et al. A streamlined workflow for automated cryo focused ion beam milling. J Struct Biol 213, 107743 (2021). 10.1016/j.jsb.2021.107743

32 Al-Amoudi, A. et al. Cryo-electron microscopy of vitreous sections. EMBO J 23, 3583–3588 (2004). 10.1038/sj.emboj.7600366

33 A, A. L.-A., Dubochet, J. & Studer, D. Amorphous solid water produced by cryosectioning of crystalline ice at 113 K. J Microsc 207, 146–153 (2002). 10.1046/j.1365-2818.2002.01051.x

34 Al-Amoudi, A., Norlen, L. P. & Dubochet, J. Cryo-electron microscopy of vitreous sections of native biological cells and tissues. J Struct Biol 148, 131–135 (2004). 10.1016/j.jsb.2004.03.010

35 Norlen, L., Al-Amoudi, A. & Dubochet, J. A cryotransmission electron microscopy study of skin barrier formation. J Invest Dermatol 120, 555–560 (2003). 10.1046/j.1523-1747.2003.12102.x

36 Han, H. M., Huebinger, J. & Grabenbauer, M. Self-pressurized rapid freezing (SPRF) as a simple fixation method for cryo-electron microscopy of vitreous sections. J Struct Biol 178, 84–87 (2012). 10.1016/j.jsb.2012.04.001

37 Studer, D., Graber, W., Al-Amoudi, A. & Eggli, P. A new approach for cryofixation by high-pressure freezing. J Microsc 203, 285–294 (2001). 10.1046/j.1365-2818.2001.00919.x

38 Moor, H. in Cryotechniques in Biological Electron Microscopy (eds Rudolf Alexander Steinbrecht & Karl Zierold) 175–191 (Springer Berlin Heidelberg, 1987).

39 Studer, D., Klein, A., Iacovache, I., Gnaegi, H. & Zuber, B. A new tool based on two micromanipulators facilitates the handling of ultrathin cryosection ribbons. J Struct Biol 185, 125–128 (2014). 10.1016/j.jsb.2013.11.005

40 Henderikx, R. J. M. et al. Ice thickness control and measurement in the VitroJet for time-efficient single particle structure determination. J Struct Biol 216, 108139 (2024). 10.1016/j.jsb.2024.108139

41 Matzelle, T. R., Gnaegi, H., Ricker, A. & Reichelt, R. Characterization of the cutting edge of glass and diamond knives for ultramicrotomy by scanning force microscopy using cantilevers with a defined tip geometry. Part II. J Microsc 209, 113–117 (2003). 10.1046/j.1365-2818.2003.01119.x

42 Gilbert, M. A. G. et al. CryoET of beta-amyloid and tau within postmortem Alzheimer’s disease brain. Nature 631, 913–919 (2024). 10.1038/s41586-024-07680-x

43 Al-Amoudi, A., Studer, D. & Dubochet, J. Cutting artefacts and cutting process in vitreous sections for cryo-electron microscopy. Journal of Structural Biology 150, 109–121 (2005). 10.1016/j.jsb.2005.01.003

44 Studer, D. & Gnaegi, H. Minimal compression of ultrathin sections with use of an oscillating diamond knife. J Microsc 197, 94–100 (2000). 10.1046/j.1365-2818.2000.00638.x

45 Al-Amoudi, A., Diez, D. C., Betts, M. J. & Frangakis, A. S. The molecular architecture of cadherins in native epidermal desmosomes. Nature 450, 832–837 (2007). 10.1038/nature05994

46 Gunkel, M. et al. Higher-order architecture of rhodopsin in intact photoreceptors and its implication for phototransduction kinetics. Structure 23, 628–638 (2015). 10.1016/j.str.2015.01.015

47 Elferich, J. et al. In situ high-resolution cryo-EM reconstructions from CEMOVIS. IUCrJ 12, 502–510 (2025). 10.1107/S2052252525005196

48 Pierson, J. et al. Improving the technique of vitreous cryo-sectioning for cryo-electron tomography: electrostatic charging for section attachment and implementation of an anti-contamination glove box. J Struct Biol 169, 219–225 (2010). 10.1016/j.jsb.2009.10.001

49 Kampjut, D., Steiner, J. & Sazanov, L. A. Cryo-EM grid optimization for membrane proteins. iScience 24, 102139 (2021). 10.1016/j.isci.2021.102139

50 Basanta, B., Hirschi, M. M., Grotjahn, D. A. & Lander, G. C. A case for glycerol as an acceptable additive for single-particle cryoEM samples. Acta Crystallogr D Struct Biol 78, 124–135 (2022). 10.1107/S2059798321012110

51 Punjani, A., Rubinstein, J. L., Fleet, D. J. & Brubaker, M. A. cryoSPARC: algorithms for rapid unsupervised cryo-EM structure determination. Nat Methods 14, 290–296 (2017). 10.1038/nmeth.4169

52 Scheres, S. H. RELION: implementation of a Bayesian approach to cryo-EM structure determination. J Struct Biol 180, 519–530 (2012). 10.1016/j.jsb.2012.09.006

53 Moriscot, C., Schoehn, G. & Housset, D. High pressure freezing and cryo-sectioning can be used for protein structure determination by electron diffraction. Ultramicroscopy 254, 113834 (2023). 10.1016/j.ultramic.2023.113834

54 Jesior, J. C. Use of low-angle diamond knives leads to improved ultrastructural preservation of ultrathin sections. Scanning Microsc Suppl 3, 147–152; discussion 152-143 (1989).

55 Richter, K. Cutting artefacts on ultrathin cryosections of biological bulk specimens. Micron 25, 297–308 (1994). 10.1016/0968-4328(94)90001-9

56 Al-Amoudi, A., Dubochet, J., Gnaegi, H., Luthi, W. & Studer, D. An oscillating cryo-knife reduces cutting-induced deformation of vitreous ultrathin sections. J Microsc 212, 26–33 (2003). 10.1046/j.1365-2818.2003.01244.x

57 Cui, Y., Chen, X., Wang, Z. & Lu, Y. Ribosome purification from Escherichia coli by ultracentrifugation. Biotechnol Notes 3, 118–123 (2022). 10.1016/j.biotno.2022.12.003

58 Asarnow, D., Palovcak, E. & Cheng, Y. UCSF pyem v0.5. Zenodo 10.5281/zenodo.3576630. (2019).

59 Zheng, S. Q. et al. MotionCor2: anisotropic correction of beam-induced motion for improved cryo-electron microscopy. Nat Methods 14, 331–332 (2017). 10.1038/nmeth.4193

60 Rohou, A. & Grigorieff, N. CTFFIND4: Fast and accurate defocus estimation from electron micrographs. J Struct Biol 192, 216–221 (2015). 10.1016/j.jsb.2015.08.008

61 Kremer, J. R., Mastronarde, D. N. & McIntosh, J. R. Computer visualization of three-dimensional image data using IMOD. J Struct Biol 116, 71–76 (1996). 10.1006/jsbi.1996.0013

62 Pettersen, E. F. et al. UCSF Chimera--a visualization system for exploratory research and analysis. J Comput Chem 25, 1605–1612 (2004). 10.1002/jcc.20084

63 Emsley, P. & Cowtan, K. Coot: model-building tools for molecular graphics. Acta Crystallogr D Biol Crystallogr 60, 2126–2132 (2004). 10.1107/S0907444904019158

64 Croll, T. I. ISOLDE: a physically realistic environment for model building into low-resolution electron-density maps. Acta Crystallogr D Struct Biol 74, 519–530 (2018). 10.1107/S2059798318002425

65 Afonine, P. V. et al. Real-space refinement in PHENIX for cryo-EM and crystallography. Acta Crystallogr D Struct Biol 74, 531–544 (2018). 10.1107/S2059798318006551

66 Goddard, T. D. et al. UCSF ChimeraX: Meeting modern challenges in visualization and analysis. Protein Sci 27, 14–25 (2018). 10.1002/pro.3235

67 Naydenova, K. & Russo, C. J. Measuring the effects of particle orientation to improve the efficiency of electron cryomicroscopy. Nat Commun 8, 629 (2017). 10.1038/s41467-017-00782-3

68 Stojkovic, V. et al. Assessment of the nucleotide modifications in the high-resolution cryo-electron microscopy structure of the Escherichia coli 50S subunit. Nucleic Acids Res 48, 2723–2732 (2020). 10.1093/nar/gkaa037

