## Supplemental file for "*In situ* structure of bacterial 50S ribosomes at 3.0 Å resolution from vitreous sections"

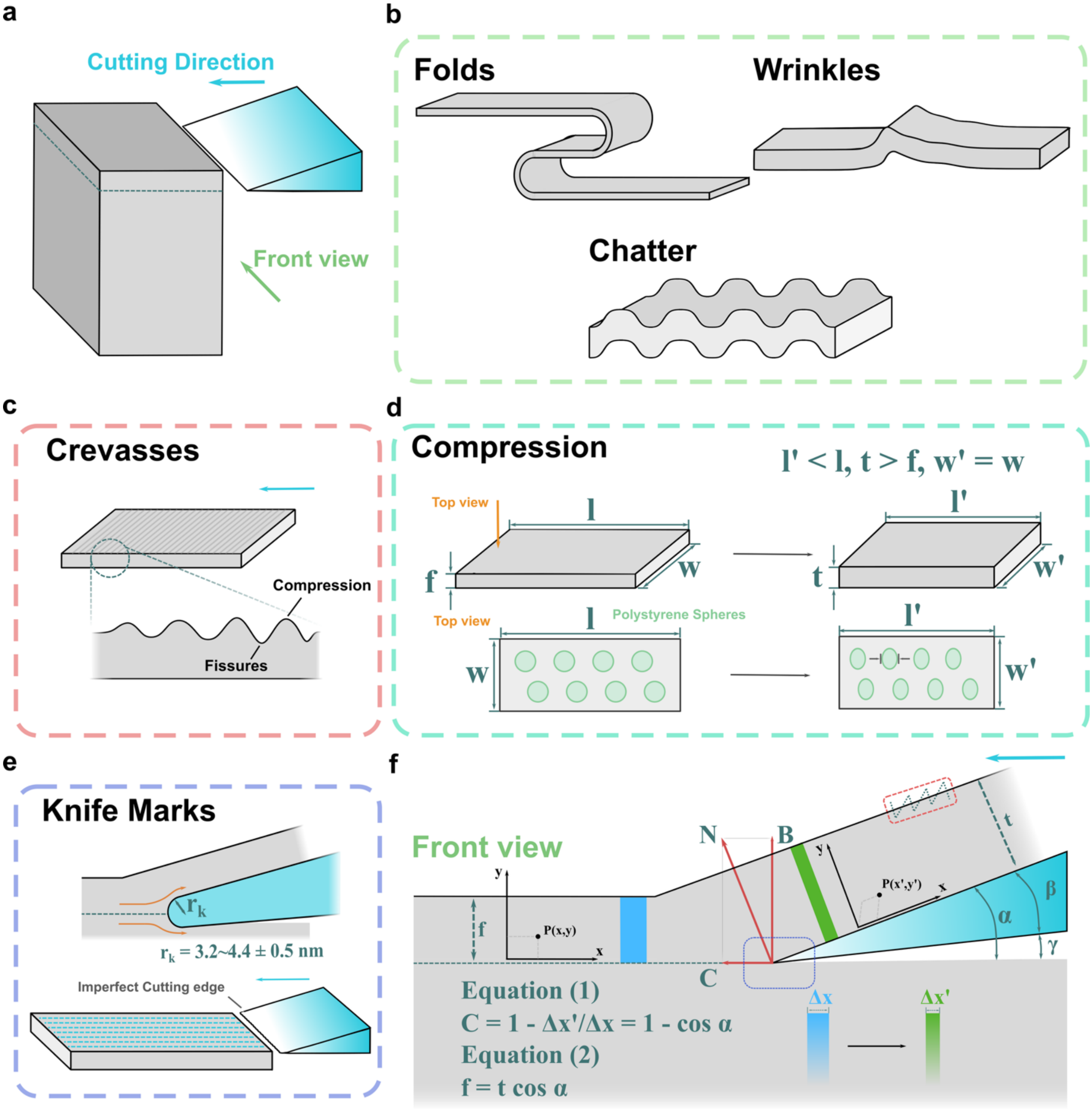


**Supplementary Figure 1: The cutting process and resulting artefacts in CEMOVIS. a,** The cutting direction is indicated. **b,** Folds, wrinkles, and chatter are artefacts that become apparent at the ultrastructural level. Folds occur when the section folds onto itself, causing parts of the section to lie on top of one another. Wrinkles appear as ripples or invaginations in the section. While folds and wrinkles are difficult to eliminate completely, they can often be avoided by collecting data from areas where they are absent. Chatter, on the other hand, is believed to result from friction at the knife surface. Newly formed sections may temporarily adhere to the knife edge, becoming immobilized. As pressure builds, the section releases abruptly, leading to periodic variations in compression, this phenomenon is known as chatter. Chatter can be completely avoided by using high-quality knives and optimizing the cutting conditions. **c,** Crevasses represent alternating regions of compressed material (dark area) at the surface and surface fissures (bright area) that become visible at high magnification. These arise from the bending of sections that are too thick. Thinner sections can bend more easily around the knife edge without significant compression of the material at the surface. In contrast, thicker sections are more difficult to bend around the knife, resulting in compression of the material near the surface, which can eventually lead to fissures caused by excessive shear strain that compromises the structural integrity of the section. Hence, crevasses can be eliminated by cutting thinner sections. **d,** Compression in the cutting direction reduces the section length $l$ into the compressed length $l'$, the feed $f$ into the final section thickness $t$, while the width $w$ remains unchanged. **e,** Knife marks depend on the radius $r_k$ of the knife edge and are unavoidable artefacts in the cutting direction, typically limited to a few nm in depth. **f,** A volume element $\Delta x$ within the section is deformed into a volume element $\Delta x'$. The compression $C$ is obtained using Equation (1). However, compression does not simply reduce the length, instead, it transforms a point $P(x, y)$ into a new point $P(x', y')$ through a more complex deformation. The compression $C$ can be measured using vitreous sections with embedded polystyrene beads, which reshape an originally spherical bead into an ellipse. The feed can be related to the section thickness after compression occurred, using Equation (2). Note: The cutting angle $\alpha$ is composed of two contributions: the knife angle $\beta$, which was set to 35°, and the clearance angle $\gamma$, which was set to 6°.

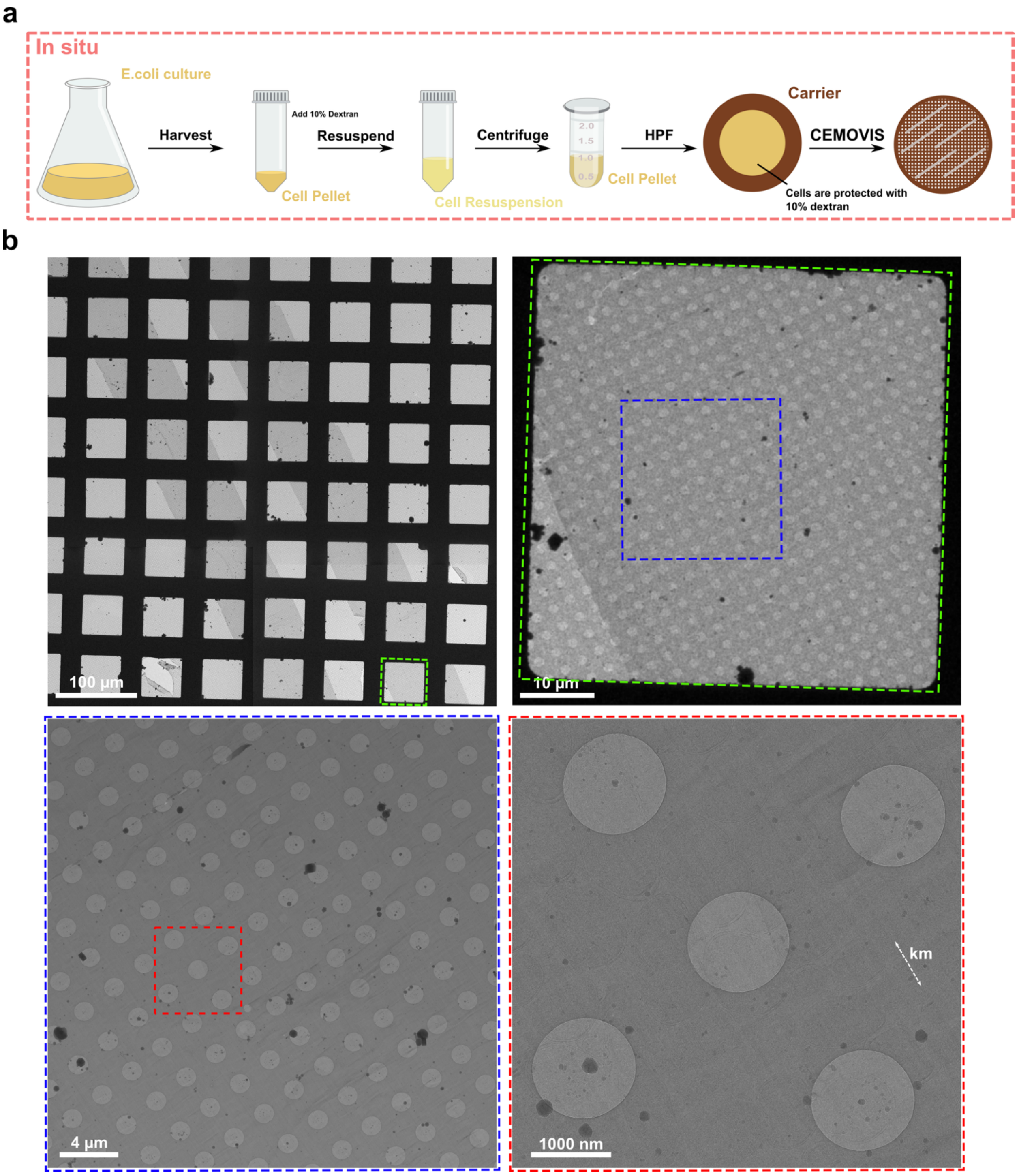


**Supplementary Figure 2: Sample preparation workflow for CEMOVIS of *E. coli* cells and assessment of section quality. a**, Preparation of *E. coli* cells for *in situ* studies.The cell culture was harvested by centrifugation to produce a pellet. Resuspension of the pellet in an equivalent volume of 20% dextran buffer yields an *E. coli* suspension with a final dextran concentration of 10%. From this suspension, 1 ml was transferred to an Eppendorf tube and centrifuged. The resulting pellet formed a dense cell paste, which was used to fill the HPF carrier. The vitrified sample was cryo-sectioned using CEMOVIS (see materials and methods). **b,** High-quality ultrathin vitreous sections from *E. coli* cells at different magnification levels. Four magnification levels are shown with scale bars. Colored boxes highlight the regions in the lower-magnification images that are shown at higher magnification in the subsequent images. Km: Knife marks.

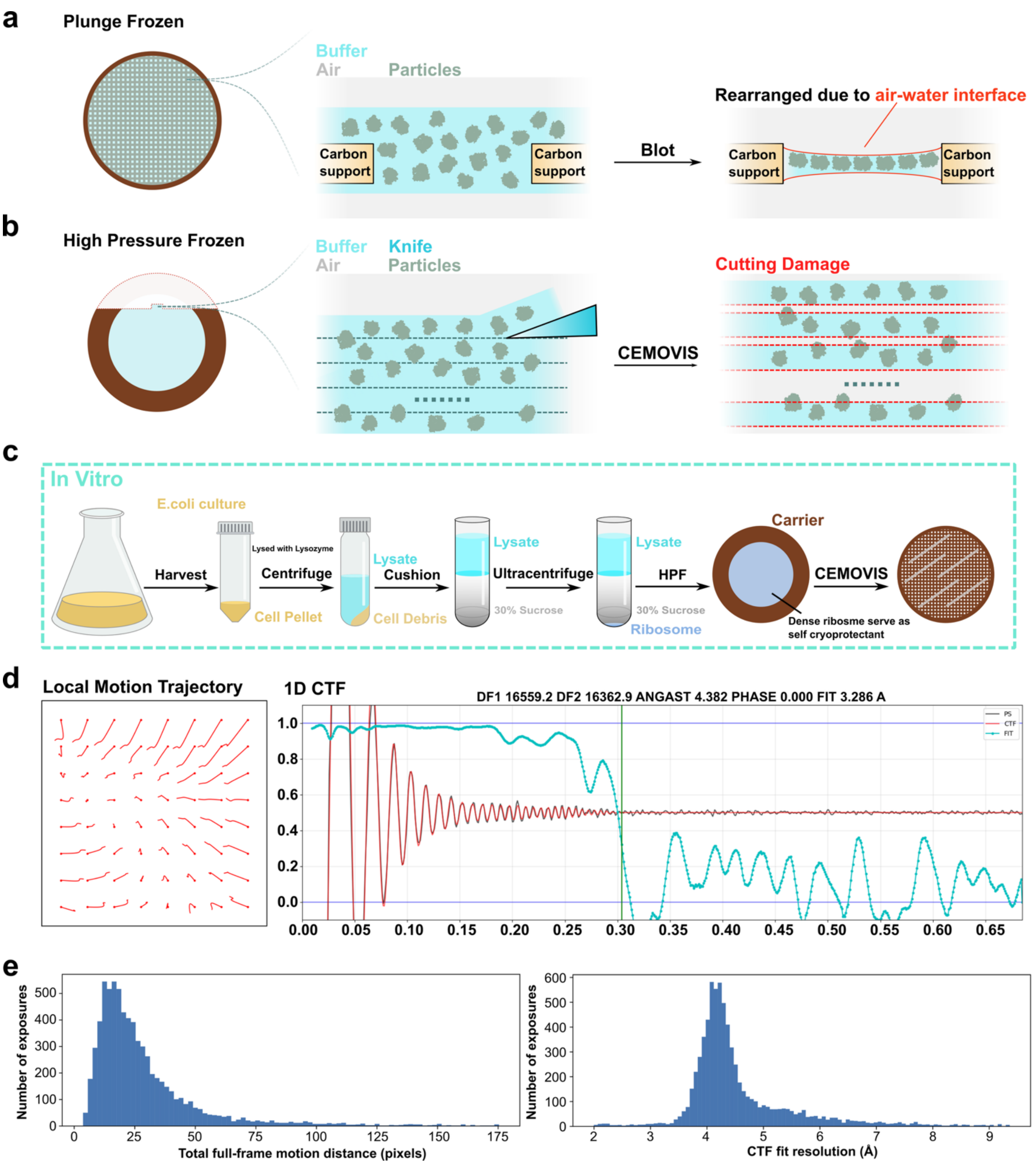


**Supplementary Figure 3: CEMOVIS applied to purified ribosomes. a,** Standard plunge-freezing of purified samples leads to preferred orientation due to interactions of particles with the air–water interface. **b,** Sections of high-pressure-frozen samples should overcome preferred orientation due to the absence of the air-water interface; however, cutting damage can affect particle integrity. **c,** Workflow for *in vitro* purified ribosomes subjected to HPF and CEMOVIS. **d,** Local motion trajectories of a representative micrograph and CTF fit versus resolution.**e,** Histograms showing the distribution of overall motion and CTF fit versus resolution across all micrographs.

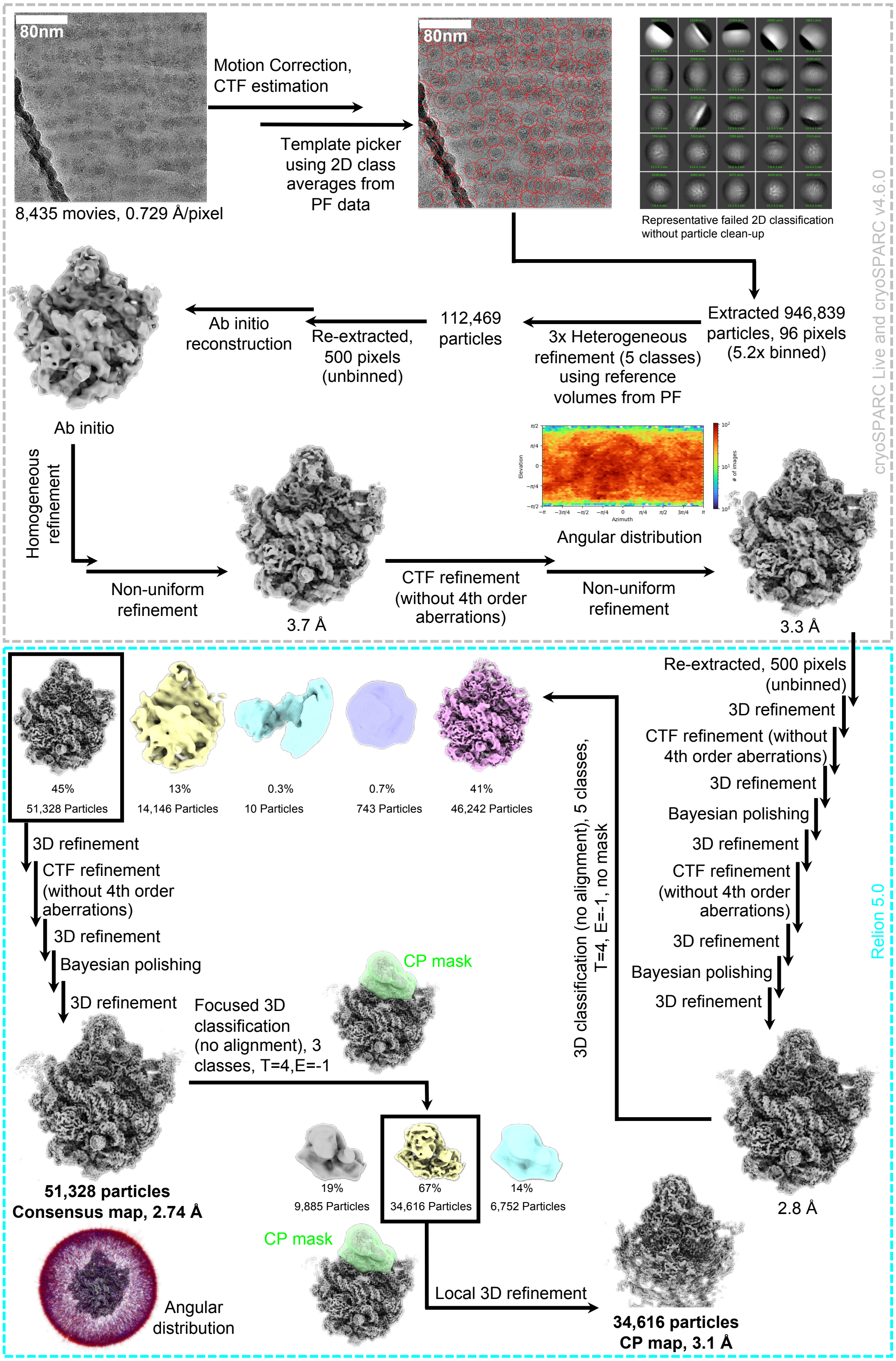

80nm
Motion Correction,
CTF estimation
Template picker
using 2D class
averages from
PF data
80nm
8,435 movies, 0.729 Å/pixel
Representative failed 2D classification
without particle clean-up
cryoSPARC Live and cryoSPARC v4.6.0
Extracted 946,839
particles, 96 pixels
(5.2x binned)
3x Heterogeneous
refinement (5 classes)
using reference
volumes from PF
112,469
particles
Re-extracted,
500 pixels
(unbinned)
Ab initio
reconstruction
Ab initio
Homogeneous
refinement
Non-uniform
refinement
3.7 Å
Angular distribution
CTF refinement
(without 4th order
aberrations)
Non-uniform
refinement
3.3 Å
Re-extracted, 500 pixels
(unbinned)
3D refinement
CTF refinement (without
4th order aberrations)
3D refinement
Bayesian polishing
3D refinement
CTF refinement
(without 4th order
aberrations)
3D refinement
Bayesian polishing
3D refinement
2.8 Å
Relion 5.0
3D classification (no alignment), 5 classes,
T=4, E=-1, no mask
45%
51,328 Particles
13%
14,146 Particles
0.3%
10 Particles
0.7%
743 Particles
41%
46,242 Particles
3D refinement
CTF refinement
(without 4th order
aberrations)
3D refinement
Bayesian polishing
3D refinement
Focused 3D
classification
(no alignment), 3
classes, T=4,E=-1
CP mask
51,328 particles
Consensus map, 2.74 Å
Angular
distribution
19%
9,885 Particles
67%
34,616 Particles
14%
6,752 Particles
CP mask
Local 3D refinement
34,616 particles
CP map, 3.1 Å

**Supplementary Figure 4: Flowchart of the data processing for purified ribosomes used in SPA of vitreous sections.** The gray box outlines all steps performed in cryoSPARC, while the turquoise box outlines the steps performed in RELION. The white scale bar in the micrograph corresponds to a length of 80 nm. Each black arrow indicates a completed job, with the job type labeled next to the arrow. Particle numbers, overall resolution, and class percentages are indicated. For the CP region, focused classification and refinement were performed using a smaller mask (colored in green) covering the entire CP region.

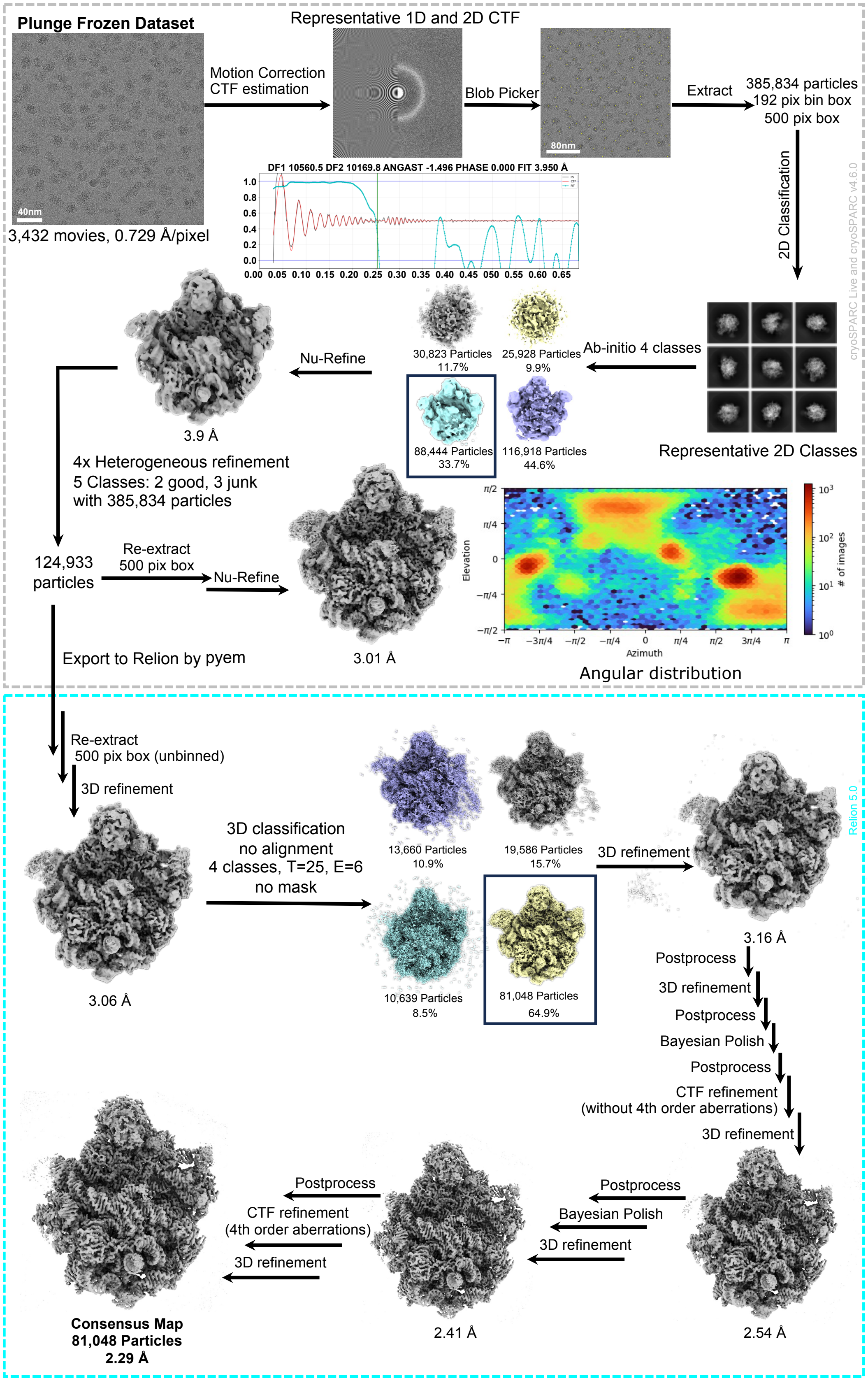

Plunge Frozen Dataset
40nm
3,432 movies, 0.729 Å/pixel
Motion Correction
CTF estimation
Representative 1D and 2D CTF
DF1 10560.5 DF2 10169.8 ANGAST -1.496 PHASE 0.000 FIT 3.950 Å
Blob Picker
80nm
Extract
385,834 particles
192 pix bin box
500 pix box
2D Classification
cryoSPARC Live and cryoSPARC v4.6.0
Representative 2D Classes
Ab-initio 4 classes
30,823 Particles
11.7%
25,928 Particles
9.9%
88,444 Particles
33.7%
116,918 Particles
44.6%
Nu-Refine
3.9 Å
4x Heterogeneous refinement
5 Classes: 2 good, 3 junk
with 385,834 particles
124,933
particles
Re-extract
500 pix box
Nu-Refine
3.01 Å
Elevation
Azimuth
### of images
Angular distribution
Export to Relion by pyem
Re-extract
500 pix box (unbinned)
3D refinement
3.06 Å
3D classification
no alignment
4 classes, T=25, E=6
no mask
13,660 Particles
10.9%
19,586 Particles
15.7%
10,639 Particles
8.5%
81,048 Particles
64.9%
3D refinement
Relion 5.0
3.16 Å
Postprocess
3D refinement
Postprocess
Bayesian Polish
Postprocess
CTF refinement
(without 4th order aberrations)
3D refinement
2.54 Å
Postprocess
Bayesian Polish
3D refinement
2.41 Å
Postprocess
CTF refinement
(4th order aberrations)
3D refinement
Consensus Map
81,048 Particles
2.29 Å

**Supplementary Figure 5: Flowchart of the data processing of plunge-frozen bacterial 50S ribosome.** The gray box outlines all steps performed in cryoSPARC, while the turquoise box outlines the steps performed in RELION. Each job type is indicated by a labeled arrow. The resolution of the reconstruction is shown beneath the maps, and the percentage of each class is provided. The angular distribution plot shows the dominant angles found in the plunge-frozen dataset. The white scale bars in the micrographs correspond to a length of 20 nm and 80nm respectively.

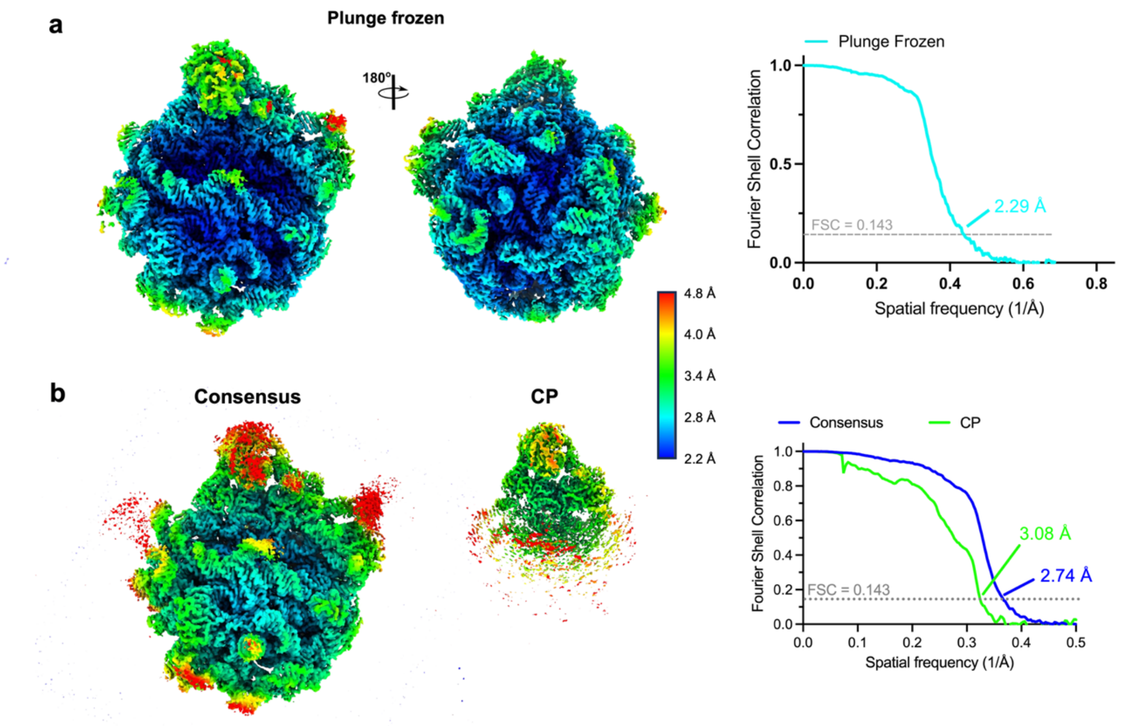


**Supplementary Figure 6: Local resolution and Fourier shell correlation (FSC) curves of purified 50S ribosomes. a**, Consensus map of plunge-frozen 50S ribosomes shown in isosurface view, colored by local resolution, with corresponding FSC curve. **b**, Consensus and CP-focused refined maps of the 50S ribosome from vitreous sections shown in isosurface representation, colored by local resolution. The right panel shows the FSC curves corresponding to the two maps on the left.

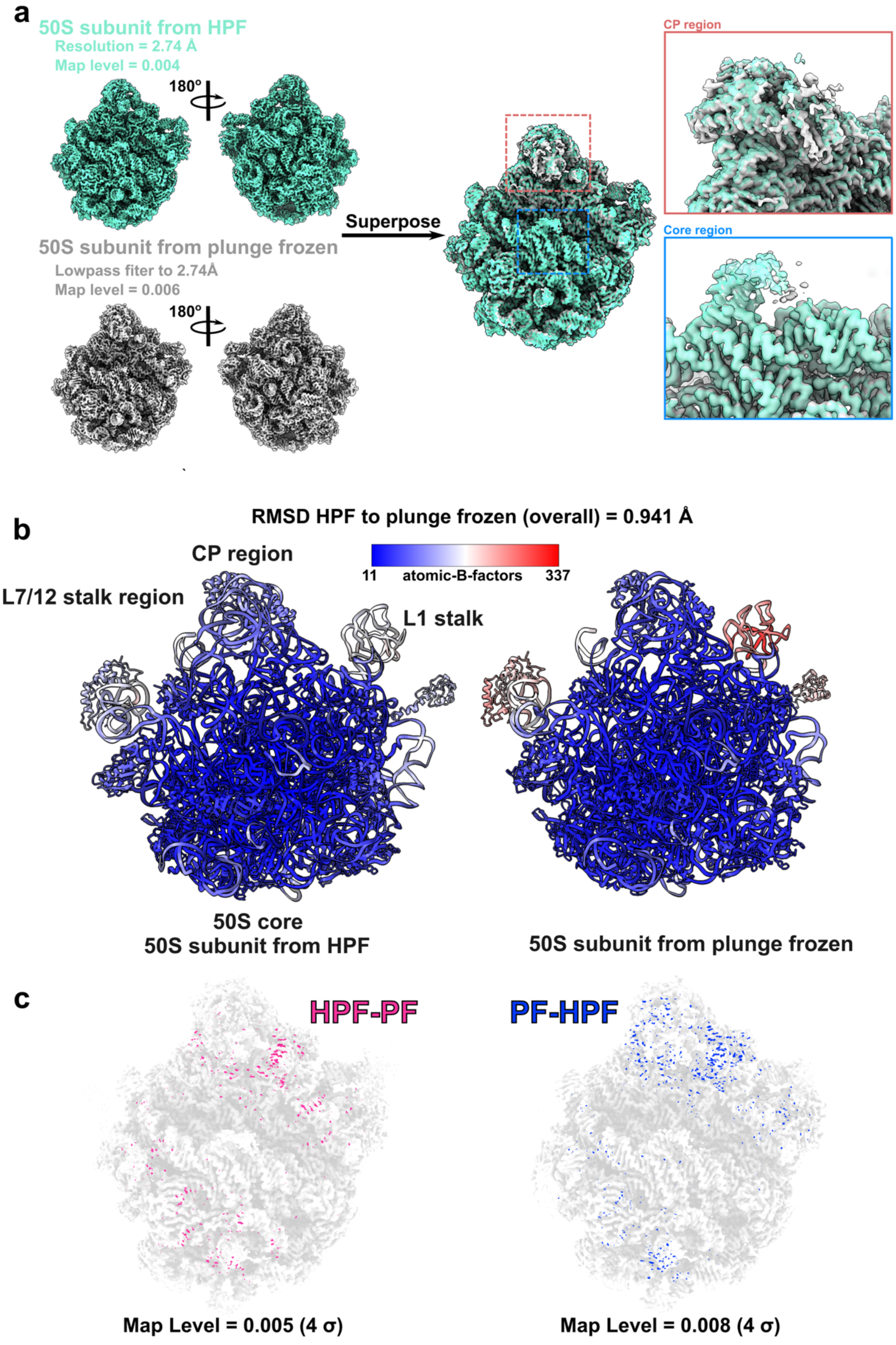

a
50S subunit from HPF
Resolution = 2.74 Å
Map level = 0.004
180°
50S subunit from plunge frozen
Lowpass fiter to 2.74Å
Map level = 0.006
180°
Superpose
CP region
Core region
b
RMSD HPF to plunge frozen (overall) = 0.941 Å
CP region
11
atomic-B-factors
337
L7/12 stalk region
L1 stalk
50S core
50S subunit from HPF
50S subunit from plunge frozen
c
HPF-PF
PF-HPF
Map Level = 0.005 (4 σ)
Map Level = 0.008 (4 σ)

**Supplementary Figure 7: Comparison of the 50S ribosomal subunit between purified CEMOVIS- and plunge-frozen-generated structures. a**, The 50S ribosome map from the CEMOVIS-generated reconstruction is shown in green, and the map from the plunge-frozen 50S ribosome is shown in gray. The superposition of both maps is displayed in the center. Close-up views on the right compare the regions indicated by the red and blue boxes, corresponding to the central protuberance (CP) and core regions, respectively. **b**, Superposed atomic models of the 50S ribosomal subunit from HPF and plunge-frozen samples are shown side by side and colored by atomic B-factor. The overall Cα RMSD between the two models is 0.941 Å. **c**, Difference maps were calculated by subtracting the plunge-frozen map from the HPF map and vice versa. Maps were aligned, low-pass filtered, and scaled before subtraction.

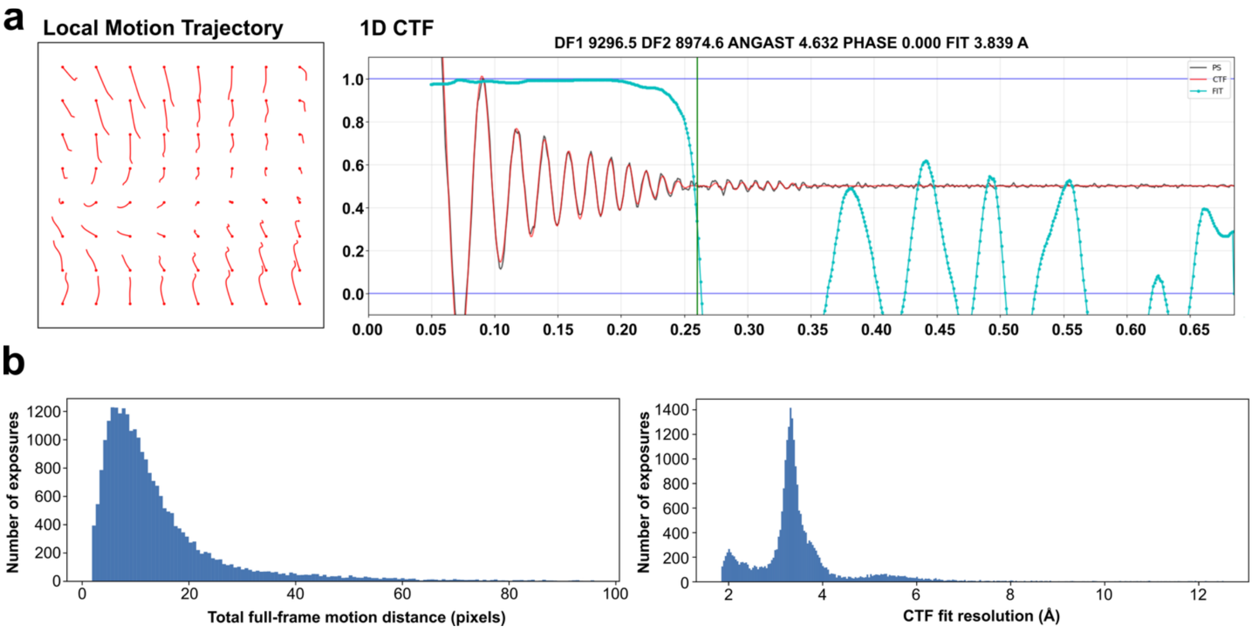


**Supplementary Figure 8: *In situ* data processing quality for the 50S ribosomal subunit. a,** Local motion trajectories and 1D CTF fit versus resolution of a representative micrograph. **b,** Histograms showing the distribution of overall motion and CTF fit versus resolution across all micrographs.

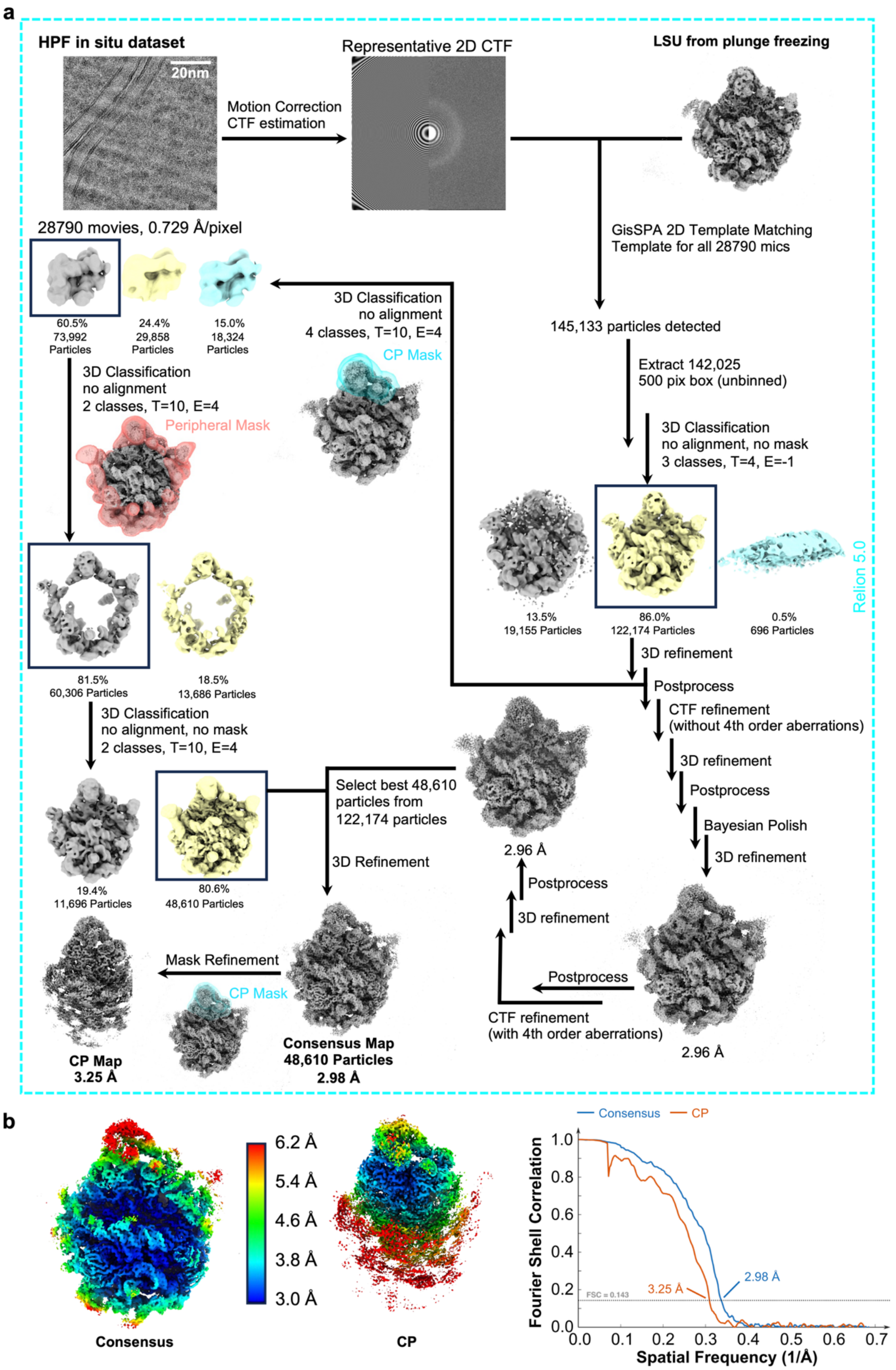
a
HPF in situ dataset
20nm
Representative 2D CTF
LSU from plunge freezing
Motion Correction
CTF estimation
28790 movies, 0.729 Å/pixel
GisSPA 2D Template Matching
Template for all 28790 mics
60.5%
73,992
Particles
24.4%
29,858
Particles
15.0%
18,324
Particles
3D Classification
no alignment
4 classes, T=10, E=4
CP Mask
145,133 particles detected
Extract 142,025
500 pix box (unbinned)
3D Classification
no alignment
2 classes, T=10, E=4
Peripheral Mask
3D Classification
no alignment, no mask
3 classes, T=4, E=-1
Relion 5.0
13.5%
19,155 Particles
86.0%
122,174 Particles
0.5%
696 Particles
81.5%
60,306 Particles
18.5%
13,686 Particles
3D refinement
Postprocess
CTF refinement
(without 4th order aberrations)
3D Classification
no alignment, no mask
2 classes, T=10, E=4
3D refinement
Postprocess
Select best 48,610
particles from
122,174 particles
Bayesian Polish
3D refinement
3D Refinement
2.96 Å
Postprocess
3D refinement
19.4%
11,696 Particles
80.6%
48,610 Particles
Mask Refinement
CP Mask
Postprocess
CTF refinement
(with 4th order aberrations)
CP Map
3.25 Å
Consensus Map
48,610 Particles
2.98 Å
2.96 Å
b
6.2 Å
5.4 Å
4.6 Å
3.8 Å
3.0 Å
Consensus
CP
Consensus
CP
Fourier Shell Correlation
1.0
0.8
0.6
0.4
0.2
0.0
FSC = 0.143
3.25 Å
2.98 Å
0.0
0.1
0.2
0.3
0.4
0.5
0.6
0.7
Spatial Frequency (1/Å)

**Supplementary Figure 9: Data processing scheme for the *in situ* 50S ribosomes using GisSPA, with local resolution and FSC curves.** Processing was carried out using RELION [1] and GisSPA [2]. Particle numbers are indicated, and each processing step is labeled with arrows and job types. For the masked classifications, the masks used to focus on specific regions are shown. The final consensus map and CP-focused map are depicted in isosurface representation, colored by local resolution, in the bottom panel along with the corresponding FSC curves. The white scale bar in the micrograph corresponds to a length of 20 nm.

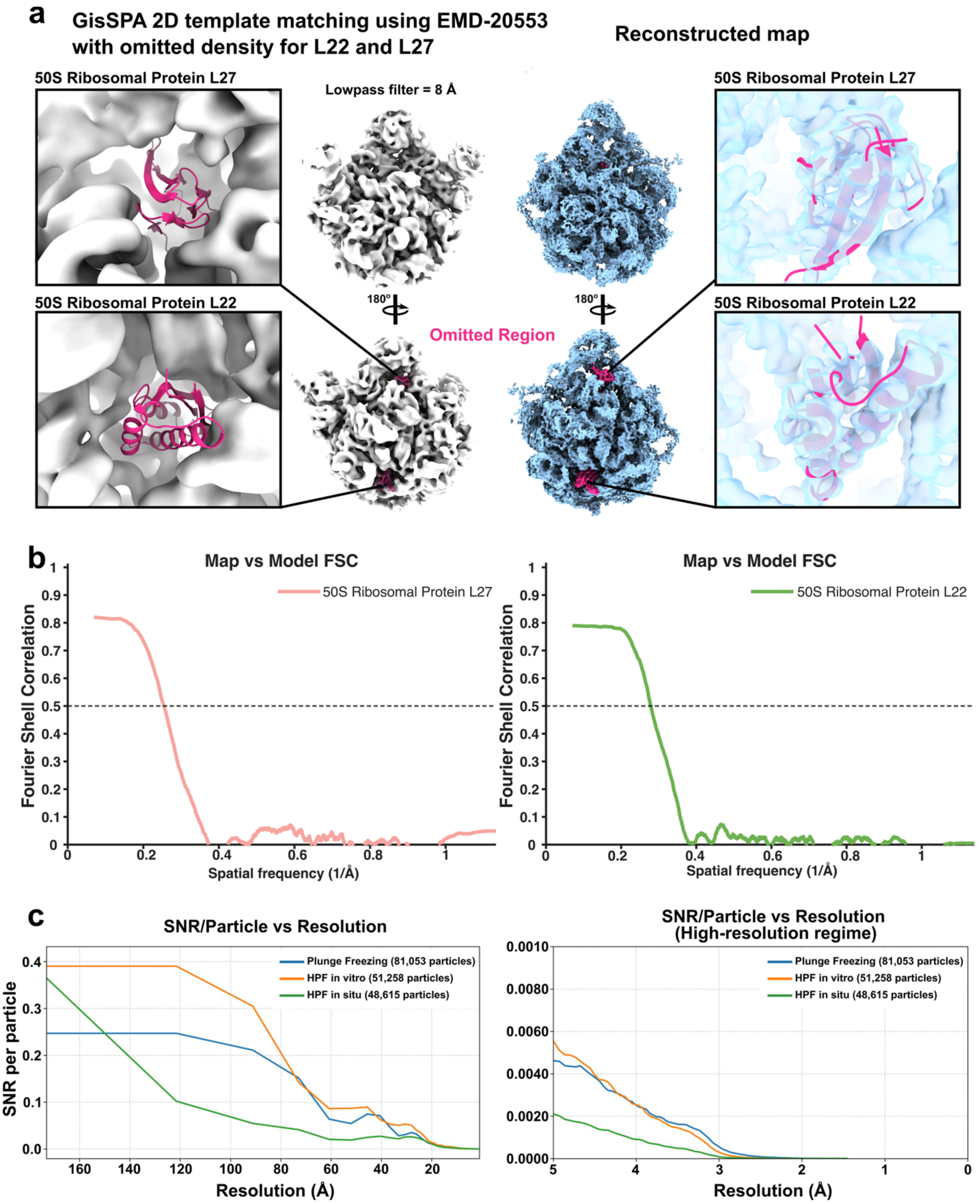


**Supplementary Figure 10: Validation of the *in situ* 50S ribosome reconstruction for template bias and SNR analysis. a,** The reference for template matching was downloaded from the PDB and low-pass filtered to 8 Å. The two subunits L22 (core region) and L27 (CP region) were omitted. Template matching enabled the reconstruction of a 3.3 Å map with well-ordered density for both subunits. **b,** Map-to-model FSC for the individual subunits. **c,** Mean per-particle SNR calculated from the half-maps and compared between plunge-frozen, *in vitro*, and *in situ* datasets.

**Supplementary Movie 1: High-pressure-frozen *E. coli* sample in a 100 µm carrier containing 20% dextran as a cryoprotectant, sectioned inside a Leica EM UC7/FC7 cryo-ultramicrotome.** The video is time-accelerated and shows the attachment of six ribbons with ~40 nm thickness within 30 minutes, corresponding to approximately 130 sections. Ribbons are attched onto 300-mesh copper Quantifoil R1.2/1.3. Sections were collected using a double micromanipulator system, and attachment to the grids was achieved by brief electrostatic charging. Prior to sectioning, the sample face was trimmed to expose vitrified material and create a cubic-like sample block; the trimming process is not shown in the video.

#### Supplementary Table 1: Success rate of CEMOVIS sample preparations.

| DATASET | RIBBONS LANDED ONTO THE GRID | GRIDS PREPARED | GOOD RIBBON QUALITY | MICROGRAPHS COLLECTED | MICROGRAPHS USED | PARTICLES EXTRACTED | PICKING METHOD |
|---|---|---|---|---|---|---|---|
| **IN VITRO** | ~60% | 3 | ~50% | ~8,435 | ~5,000 (~63%) | 946839 | Blob picking (cryosparc) |
| **IN SITU** | ~80% | 4 | ~65% | 28,790 | ~14,000 (~50%) | 142025 | GisSPA |

1 Scheres, S. H. RELION: implementation of a Bayesian approach to cryo-EM structure determination. *J Struct Biol* **180**, 519-530 (2012). https://doi.org:10.1016/j.jsb.2012.09.006
2 Cheng, J. *et al.* Determining protein structures in cellular lamella at pseudo-atomic resolution by GisSPA. *Nat Commun* **14**, 1282 (2023). https://doi.org:10.1038/s41467-023-36175-y
